# A Neutrophil Defensin–VDAC1 Axis Drives Steatotic Liver Disease

**DOI:** 10.64898/2026.09.19.752841

**Authors:** Jue Zhang, Gan Luo, Gang Wang, Jie Rao, Xiao Li, Wuyuan Lu

**Affiliations:** Key Laboratory of Medical Molecular Virology (MOE/NHC/CAMS), Shanghai Institute of Infectious Disease and Biosecurity, School of Basic Medical Sciences, Fudan University Shanghai Medical College, Shanghai 200032, China; Department of General Surgery, Xijing Hospital, Air Force Medical University, Xi’an 710032, China

## Abstract

Neutrophils infiltrate the liver as metabolic dysfunction-associated steatotic liver disease (MASLD) progresses from simple steatosis to steatohepatitis, but whether neutrophil effectors merely mark this transition or cause it is unknown. Here we show that human neutrophil peptides (HNPs), or α-defensins, are drivers rather than bystanders of disease. In patients surgically treated for hepatic hemangioma who displayed overt MASLD, the abundance of HNPs in their liver biopsy samples localized to diseased tissue correlates with the histological severity of steatosis. In vitro, HNPs enter hepatocytes to induce multimerization of voltage-dependent anion channel 1 (VDAC1) in the mitochondrial outer membrane, causing mitochondrial dysfunction and lipid accumulation. Under lipid overload, but not on a standard diet, HNP1-transgenic mice develop increased adiposity and steatohepatitis, which is reversed by an orally available prodrug that inhibits HNP-induced VDAC1 multimerization. These findings link neutrophil innate immunity to hepatocyte mitochondrial dysfunction, recast host-protective HNPs as pathogenic effectors when hepatocyte mitochondria are lipid-burdened, and establish the HNP–VDAC1 axis as a therapeutic target in MASLD.

## INTRODUCTION

Metabolic dysfunction-associated steatotic liver disease (MASLD), formerly known as nonalcoholic fatty liver disease (NAFLD), is the most common chronic liver disease in the world^1,2^, affecting up to 30% of the global population^3,4^. It defines a spectrum of abnormalities ranging from benign steatosis (MASL) to pathological steatohepatitis (MASH), with the latter characterized by inflammatory infiltration, mitochondrial dysfunction, endoplasmic reticulum stress and fibrosis^5–7^ carrying a substantially elevated risk of cirrhosis and hepatocellular carcinoma^8,9^. Why and how simple steatosis progresses to pathogenic steatohepatitis remains the central unanswered question in the field of liver metabolism, and the principal obstacle to therapeutic intervention^10^.

Since hepatic inflammation and the resulting cellular and tissue injury are the defining hallmarks of the progression of MASL to MASH^11,12^, inflammatory and immune signaling pathways have been studied extensively for their contribution to MASLD pathogenesis^6^. Neutrophils, however, have received far less attention than liver-resident macrophages^13^ — a striking omission given that they account for 50–70% of circulating leukocytes and carry the most potent antimicrobial and cytotoxic arsenal of any granulocyte^14^. Although scarce in a healthy liver, neutrophils as the first responders of the immune system are rapidly recruited from circulation in massive numbers into tissue exposed to inflammatory conditions such as MASLD^15,16^. Evidence from animal models and clinical cohorts now suggests that this influx exacerbates inflammation and contributes to MASLD progression^6,17,18^, but the effectors through which neutrophils act, and their cellular targets in the liver, remain poorly defined.

Tissue-infiltrating neutrophils are short-lived and, upon completion of their surveillance and defense mission, mostly undergo programmed cell death to maintain immune homeostasis^19^. Among their most abundant effectors are the cationic, Cys-rich α-defensins 1–4, or human neutrophil peptides 1–4 (HNPs 1–4), densely packed in azurophilic granules^20–23^ and produced by adult humans at an estimated rate of 10–15 mg/kg body weight per day^24–27^. Their canonical function is direct microbicidal killing within the phagosome^28,29^, but what becomes of HNPs disseminated into tissue by dying neutrophils, and what they do once microbial clearance is complete, remain obscure. A growing body of work shows that HNPs and their enteric counterpart human defensin 5 (HD5) possess non-microbicidal activities well beyond host defense, acting as a “double-edged sword” that can enable rather than restrain pathogenesis under certain biological settings^30–33^.

Existing reports on defensins in liver disease appear contradictory. Elevated HNP1-3 staining in advanced MASLD/MASH fibrosis correlated with histological severity in one retrospective study of human biopsies^34^, and HNP1 stimulated hepatic stellate cell proliferation to promote alcohol- or CDAA diet-induced fibrosis while exerting little influence on steatosis^35,36^. By contrast, long-term expression of neutrophil α-defensins in mice maintained on a standard chow diet was associated with increased fat utilization and reduced hepatic steatosis^37^, and HNP1 limited hypercholesterolemia-induced atherosclerosis by facilitating hepatic LDL clearance^38^. Are HNPs merely passive markers, deposited in proportion to the neutrophil infiltration that accompanies inflamed tissue? Or are they drivers, acting on hepatocytes to worsen the very disease with which they correlate?

Any such injury would most plausibly converge on hepatocyte mitochondria, the principal site of fatty acid disposal. Mitochondrial failure — impaired β-oxidation, excess reactive oxygen species and release of mitochondrial contents into the cytosol — sustains both steatosis and inflammation^39^. Voltage-dependent anion channel 1 (VDAC1), the most abundant protein of the mitochondrial outer membrane, sits at this junction as the main conduit for metabolites, nucleotides and Ca^2+^ between cytosol and mitochondrion^40–43^. Under stress, VDAC1 self-associates into higher-order oligomers that permeabilize the outer membrane and compromise mitochondrial function^40^, causing metabolic and degenerative disease^40,41,44,45^. Notably, HNPs have a strong intrinsic propensity to self-associate and a known ability to cluster target proteins^46,47^, and are, thus, plausible candidates to nucleate VDAC1 oligomerization, provided they can enter hepatocytes.

Here we ask whether hepatic HNPs are drivers of MASLD severity or markers of it. Using human liver biopsy samples from patients surgically treated for hepatic hemangioma, we demonstrate that the amount of HNPs in the liver strongly correlates with the severity of hepatic steatosis. We show that HNPs are taken up by hepatocytes, that they induce VDAC1 multimerization and consequent mitochondrial dysfunction, that HNP1-transgenic mice develop steatohepatitis and increased adiposity under a high-fat diet but not on chow, and that this phenotype is reversed by an orally available prodrug targeting HNP-induced VDAC1 multimerization. Together these results define a neutrophil defensin–VDAC1 axis as a driver of steatotic liver disease in which HNPs from infiltrating neutrophils become pathogenic when hepatocyte mitochondria are already burdened by lipid overload.

## RESULTS

### Elevated levels of HNPs in the liver strongly correlate with the severity of hepatic steatosis in clinical patients

HNPs 1-3 differing from each other by a single amino acid residue at their N-terminus constitute the bulk of α-defensins produced by neutrophils^22,23^, whereas their “distal cousin” HNP4 is expressed only in a minuscule quantity (**Figure S1**)^48^. To assess the clinical relevance of HNPs 1-3 to hepatic steatosis, we collected a total of 29 human liver biopsy samples from surgically treated patients with hepatic hemangioma, among whom 21 displayed overt steatoses to various degrees. We classified these tissue samples into three groups on the basis of the percentage of positive oil red O (ORO)-staining area as a standard metric for the severity of hepatic steatosis as described^49,50^: non-steatosis (<5%, n=8), mild steatosis (5-33%, n=14), and severe steatosis (>33%, n=7). Shown in **Figure S2a** are representative patients’ tissue samples histopathologically stained with H&E, ORO, Masson’s trichrome, and Sirius red. As expected, expanded hepatic steatosis was accompanied by increasingly deteriorative histopathological changes in the liver such as ballooned and macrovesicular hepatocytes (**Figure S2a-b**), which were further validated by elevated NAFLD activity scores (NAS) and worsened fibrosis resulting from steatosis (**Figure S2c-d**). Using Sirius red staining, different types of collagens in liver tissues were visualized under polarized light (**Figure S2a**), revealing a gradual elevation of type IV collagen (colored in yellow) associated with initial fibrogenesis in parallel with lipid deposition (**Figure S2b** and **S2e**). Meanwhile, liver tissues from the group of severe steatosis exhibited a greater deposition of type I collagen (colored in red) than those from the other groups (**Figure S2e**), predictive of the emergence of cirrhosis.

Strikingly, HNPs 1-3 levels in the liver tissues with severe steatosis were significantly higher than those in the mild steatosis or non-steatosis group, as measured by immunohistochemical (IHC) staining coupled with semi-quantitative analysis (**Figure 1a-b**). In fact, IHC scoring showed that 57.1% and 42.9% of samples from the severe steatosis group were strongly and moderately positive, respectively, whereas the mild steatosis group contained no strongly positive samples compared with 100% weakly positive samples from the non-steatosis group (**Figure 1b and Figure S2f**). Importantly, hepatic HNPs 1-3 levels strongly correlated with areas of steatosis (*P* < 0.0001, R^2^ = 0.8579), NAS (*P* < 0.0001, R^2^ = 0.6611), and areas of fibrosis (*P* < 0.0001, R^2^ = 0.6038) (**Figure 1c-d and Figure S2g**), providing statistical evidence for a potential clinical association of HNPs 1-3 with development and progression of steatotic liver disease.

**Figure 1.**
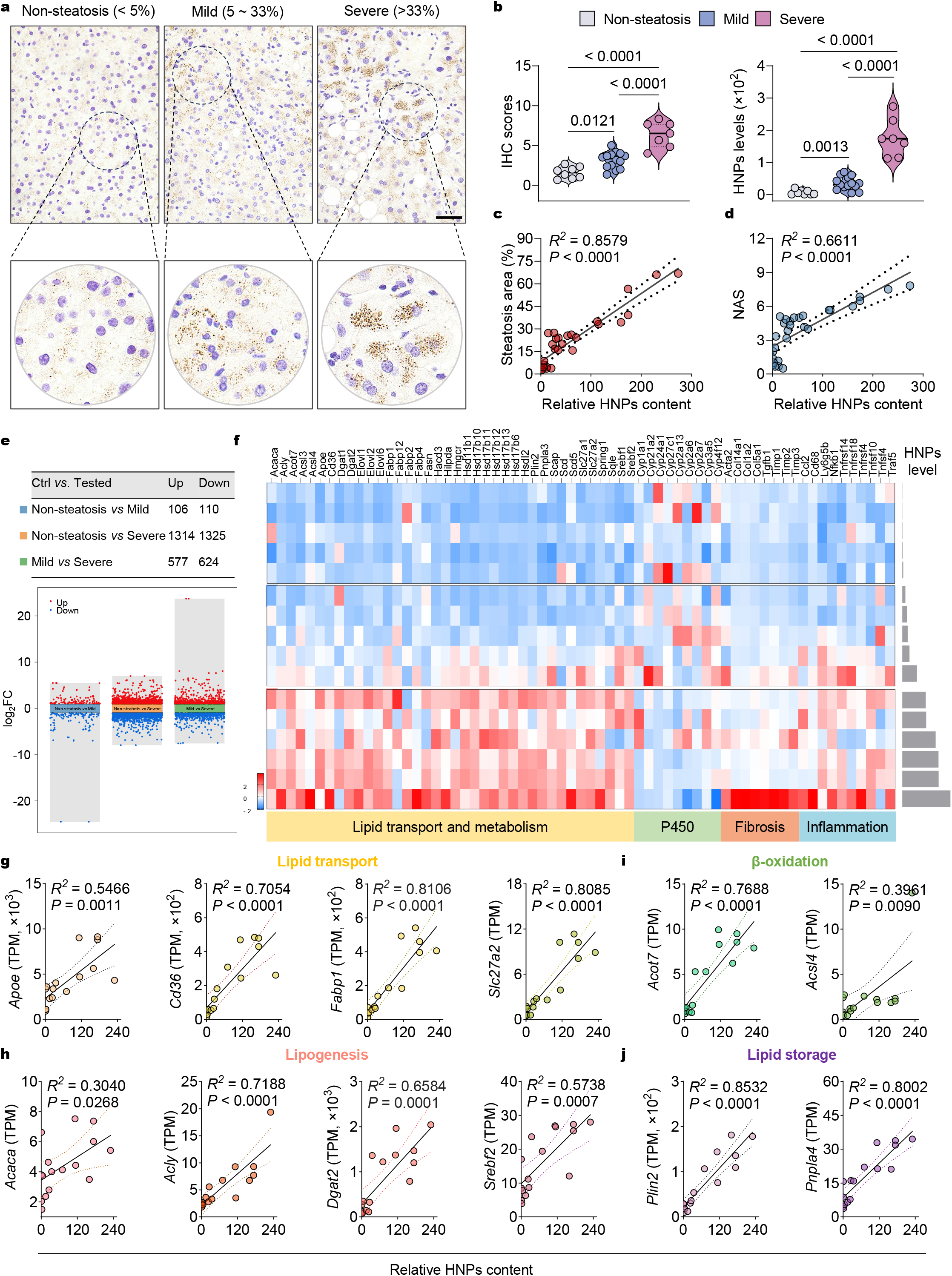
Increased infiltration of HNP1 in the liver correlates with hepatic steatosis in clinical patients. (**a**) Representative IHC images illustrating the levels and distribution of HNPs in human liver sections across different severities of hepatic steatosis. Scale bar: 100 μm. (**b**) IHC scoring of liver sections from patients without hepatic steatosis (*n* = 8) or with mild (*n* = 14) and severe steatosis (*n* = 7), and semi-quantitative analysis of HNPs levels normalized from IHC positive areas in these samples. (**c-d**) Correlation analysis of hepatic HNPs levels with steatosis area (**c**) and with NAFLD activity scores (**d**). (**e**) Multi-volcano plots illustrating the differentially expressed genes (DEGs) in clinical samples from minimal, mild, and severe hepatic steatosis groups. (**f**) RNA-seq profiles showing expression changes of genes related to lipid transport and metabolism, P450, Fibrosis, and inflammation in sixteen human liver samples, with relative HNPs levels overlaid as bar plots. (**g-j**) Scatterplots showing the correlation between the relative HNPs content and expression of genes related to lipid transport (**g**), lipogenesis (**h**), β-oxidation (**i**), and lipid storage (**j**). Coefficients of determination (R^2^) and *P* values are shown. Statistical significance was calculated by one-way analysis of variance with the Bonferroni correction for multiple comparisons.

Further transcriptomic analysis revealed significant genetic alterations associated with hepatic steatosis as its severity progressively increased the number of differentially expressed genes (DEGs) from as few as 216 between clinical samples from the non-steatosis and mild steatosis groups to as many as 2639 from the non-steatosis and severe steatosis groups (**Figure 1e**). In the severe steatosis group, numerous DEGs enriched in lipid transport and metabolism pathways along with the genes involved in fibrosis and inflammation were significantly upregulated, while P450-related genes were downregulated, indicative of apparent liver injury (**Figure 1f**). Of note, the expression levels of these upregulated genes were generally correlated with HNPs levels, as exemplified by *Apoe*, *Cd36*, *Fbp1*, and *Slc27a2* genes involved in lipid transport (**Figure 1g**), *Acaca*, *Acly*, *Dgat2*, and *Srebf2* genes related to lipogenesis (**Figure 1h**), *Acot7* and Acsl4 genes associated with β-oxidation (**Figure 1i**); and *Plin2* and *Pnpla4* genes relevant to lipid storage (**Figure 1j**). Taken together, our findings from clinical patients’ biopsy samples establish a strong correlation between the severity of hepatic steatosis and elevated levels of HNPs in the liver likely derived from increased infiltrating neutrophils under chronic inflammatory conditions.

### HNP1 transgenic mice fed with high-fat diet are prone to develop MASLD

Mice do not express defensins in neutrophils^51^. For phenotypic verification in an animal model, HNP1 transgenic (*HNP1^TG^*) mice in a C57BL/6J background were genetically constructed by CRISPR-Cas9 using neutrophil elastase gene as a promoter. Upon confirmation of a successful knock-in of *DEFA1* encoding HNP1 (*homo sapiens*, NCBI ID: 1667) (**Figure S3**), male *HNP1^TG^* mice at six weeks old, along with wild-type C57BL/6J mice in a control group, were fed with high-fat diet (HFD) for 12 weeks (**Figure 2a**). Despite nearly identical food intake (**Figure S4a**), *HNP1^TG^* mice became substantially more obese than their wild-type (WT) counterparts, as evidenced by an increased growth rate (**Figure 2b** and **Figure S4b**), resulting in significantly enlarged body and liver volumes (**Figure 2c**). In fact, the transgenic mice were on average 14.7%, 18.1% and 10.6% heavier than WT controls at 4, 8 and 12 weeks of HFD feeding (**Figure 2d**), respectively, suggesting HNP1-induced systemic fat accumulation and metabolic disturbance under HFD feeding. In sharp contrast, we failed to observe this phenotypic difference between *HNP1^TG^* and WT mice under control-fat diet (CFD) feeding (**Figure S5a-d**).

**Figure 2.**
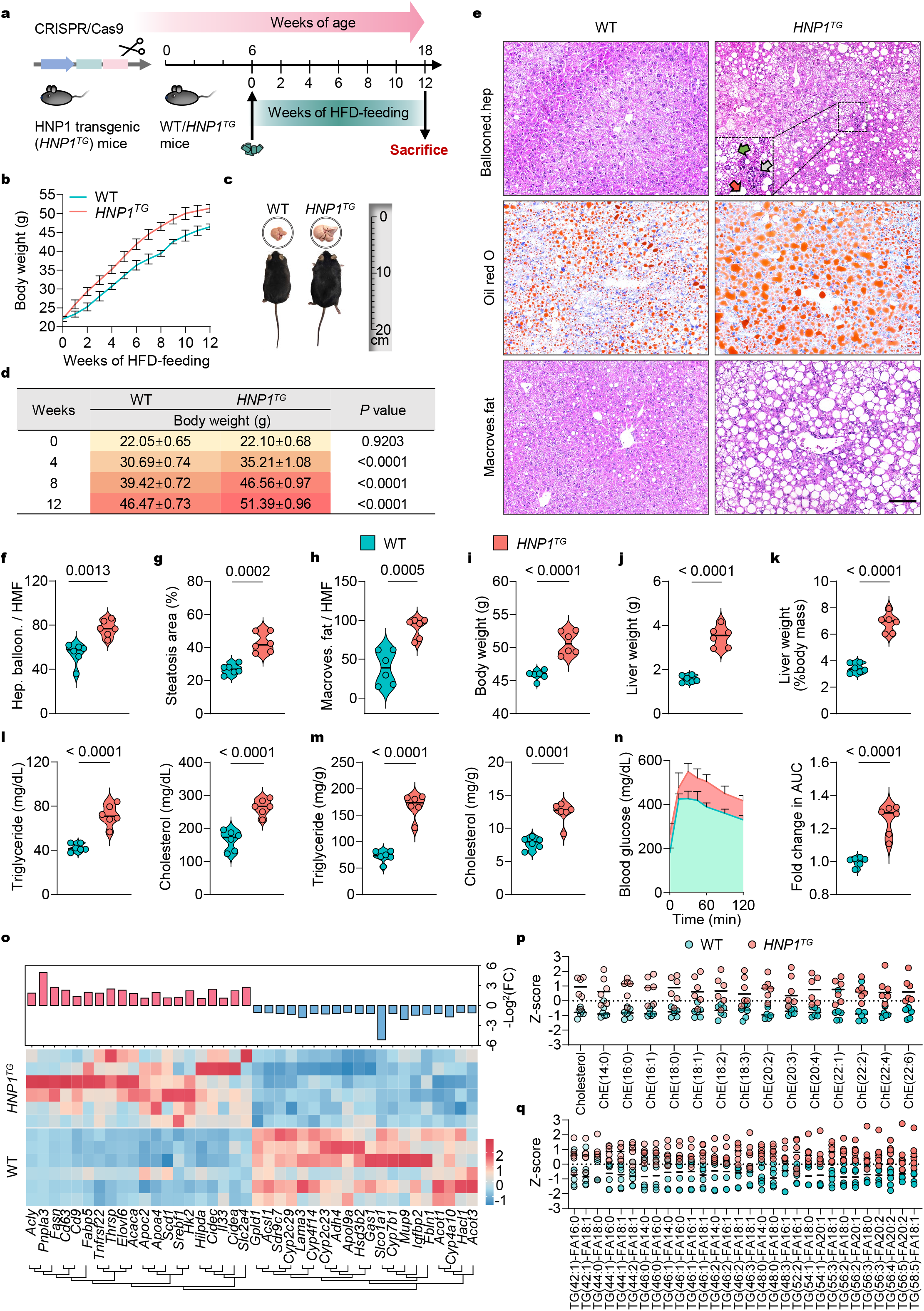
Elevated susceptibility to MASLD in HNP1 transgenic mice. (**a**) Experimental design for evaluating the role of HNP1 in MASLD progression. Six-week-old male WT and *HNP1^TG^* C57BL/6J mice, weighed 21-23 g, were fed a HFD for 12 weeks to induce MASLD, after which they were sacrificed and tissue samples collected for in-depth analysis. (**b**) Body weights of WT and *HNP1^TG^* mice measured weekly during HFD feeding (*n* = 10 mice per group). (**c**) Representative photographs of WT and *HNP1^TG^* mice (bottom) and their livers (top) following 12 weeks of HFD feeding. (**d**) Summary of body weights and statistical comparisons between WT and *HNP1^TG^* mice at 0, 4, 8, and 12 weeks. (**e**) Representative images of H&E- and ORO-stained mouse liver sections. Gray, green and red arrows indicate inflammatory infiltration, hepatocyte ballooning (Ballooned.hep) or macrovesicular fat (Macroves.fat), respectively. Scale bar: 200 μm. (**f-h**) Quantification of ballooned hepatocytes (**f**), percentage of ORO-positive area (**g**), and hepatocytes with macrovesicular fat (**h**) per high magnification field (HMF) using ImageJ, based on three fields per section (*n* = 6 mice per group). (**i-k**) Body weights (**i**), liver weights (**j**), and liver weight/body weight ratios (**k**) of WT and *HNP1^TG^* mice after 12 weeks of HFD feeding (*n* = 6 mice per group). (**l-m**) Levels of triglyceride and cholesterol in serum (**l**) and livers (**m**) from mice after 12 weeks of HFD feeding (*n* = 6 mice per group). (**n**) Glucose tolerance test of WT and *HNP1^TG^* mice following HFD feeding for 12 weeks (left panel), with fold changes in the area under curve (AUC) of blood glucose (*n* = 6 mice per group) (right panel). (**o**) Heatmap showing expression patterns of DEGs related to lipid metabolism in livers of WT and *HNP1^TG^* mice (*n* = 6 mice per group). The color key indicates the expression levels. (**p-q**) Metabolomic analysis of cholesterol-derived (**p**) and triglyceride-derived (**q**) lipid species levels in livers of WT and *HNP1^TG^* mice (*n* = 6 mice per group). All data are shown as mean ± SD. Statistical significance was analyzed by an unpaired, 2-tailed Student’s *t* test.

Mice were sacrificed at the 12^th^ week after overnight fasting for liver and epididymal fat (EAT) pads collection for extensive biochemical and histopathological analysis. As shown in **Figure 2e**, liver tissues from transgenic mice showed, to a far greater degree, hepatocyte ballooning (green arrow), macrovesicular fat (red arrow), and an inflammatory infiltrate (gray arrow) than those from WT controls, consistent with semi-quantitative analysis of these pathological indicators of hepatic steatosis (**Figure 2f-h**). In line with these findings, both terminal body and liver weights ticked up considerably with the pathological changes (**Figure 2i-j**), yielding an at least two-fold higher liver weight index (liver-to-body weight ratio) for transgenic mice over WT controls (**Figure 2k**). Interestingly, EAT pad weights from *HNP1^TG^* mice were reduced by approximately 25% compared to those of WT animals (**Figure S4c-d**). Further, *HNP1^TG^* mice exhibited enlarged adipocytes and the appearance of typical crown-like structures (CLS) in EAT pads relative to WT mice (**Figure S4e-g**), indicative of systemic lipid remodeling accompanied by aggravated chronic inflammation in adipose tissues.

From a clinical perspective, all *HNP1^TG^* mice progressed to the stage of MASH (NAS > 5) as opposed to none of the WT mice (**Figure S4h**). In this regard, numerous fiber bundles, particularly in the interlobular space, were observed in the livers of *HNP1^TG^* mice, occupying approximately 9% of the microscopic field area, whereas such fibers were nearly undetectable in WT mice (**Figure S4i-j**). Consequently, *HNP1^TG^* mice exhibited significantly higher levels of triglyceride and cholesterol in both serum and liver compared to WT controls (**Figure 2l-m**), concomitant with overt liver injury (**Figure S4k-l**). Of note, HNP1 knock-in led to a 50% increase in fasting blood glucose (**Figure S4m**). Further, a glucose tolerance test (GTT) revealed a reduced capacity for blood glucose regulation in *HNP1^TG^* mice after 12 weeks of HFD feeding (**Figure 2n**). Accordingly, insulin sensitivity was also diminished in transgenic mice compared to WT mice (**Figure S4n-o**), suggesting the emergence of insulin resistance. As expected, none of the above indicators showed significant differences between *HNP1^TG^* and WT mice fed with CFD for 12 weeks (**Figure S5e-t**).

We further retrieved transcriptomics and metabolomics on the livers from WT and *HNP1^TG^* mice fed with 12-week HFD. Generally, the samples from *HNP1^TG^* mice projected a well-separated cluster distinct from WT samples in the two-dimensional space of a principal component analysis (PCA) of both omics datasets (**Figure S6**), suggesting a significant difference between the two genotypes of mice in response to HFD feeding. As expected, in addition to transcriptional changes in genes highly relevant to lipid metabolism (**Figure 2o**), we observed elevated levels of cholesterol and triglyceride as well as their long-chain species, signifying insufficient L-carnitine shuttle of long-chain fatty acids from the cytosol to the mitochondria in *HNP1^TG^* mice (**Figure 2p-q**). Collectively, these results unequivocally demonstrate a genetic predisposition of *HNP1^TG^* mice to MASLD under HFD, implicating the neutrophil α-defensin(s) as an essential enabler of hepatic steatosis in humans.

### HNP1 induces mitochondrial dysfunction to facilitate lipid accumulation in hepatocytes

To elucidate the detrimental role of HNP1 in hepatic steatosis *in vitro*, a cellular lipotoxicity model was established with mouse hepatocyte cell line BNL CL.2, primary mouse hepatocytes or human hepatocyte cell line THLE-2 in the presence of 0.5 mM palmitic acid as a free fatty acid (FFA) (**Figure 3a**). Consistent with the *in vivo* observations, all three types of hepatocytes accumulated substantially more lipid upon treatment with 4 μM HNP1, but not with a linear HNP1 peptide (L-HNP1) at the same concentration (**Figure 3b-c**), suggesting that the HNP1-induced increase in lipid deposition is structure-dependent.

**Figure 3.**
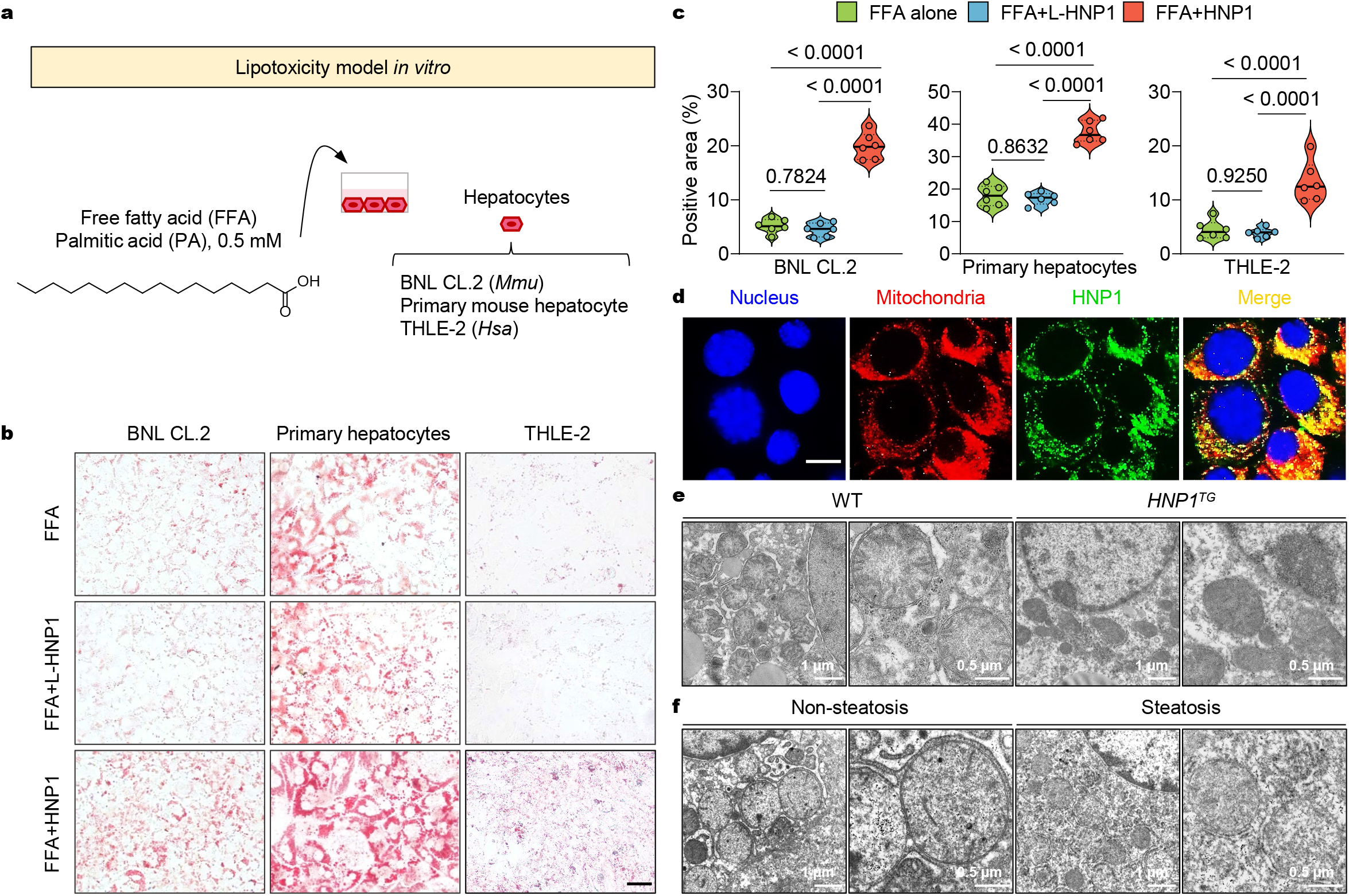
HNP1 induces mitochondrial dysfunction to facilitate hepatocyte steatosis. (**a**) Experimental diagram of the *in vitro* lipotoxicity model established in three types of hepatocytes treated with 0.5 mM palmitic acid. (**b**) Representative images of ORO-stained hepatocytes treated with 0.5 mM palmitic acid in the presence or absence of 4 μM HNP1 or linear HNP1. Scale bar: 100 μm. (**c**) Semi-quantitative analysis of lipid accumulation in hepatocytes based on the percentages of ORO-positive area (*n* = 6 biological replicates). Data are shown as mean ± SD. Statistical significance was analyzed by one-way analysis of variance with the Bonferroni correction for multiple comparisons. (**d**) Fluorescent images of mouse hepatocyte cell line BNL CL.2 incubated with 4 μM FAM-labelled HNP1, followed by mitochondria staining with MitoTracker Red. Scale bar: 25 μm. (**e**) Representative TEM images of hepatocellular mitochondria in livers of WT and *HNP1^TG^* mice after 12 weeks of HFD feeding. (**f**) Representative TEM images of hepatocellular mitochondria in livers isolated from clinical patients with or without hepatic steatosis.

To further decipher the molecular mechanisms underlying HNP1-induced lipid accumulation, RNA-seq was performed on primary mouse hepatocytes in the presence or absence of 4 μM HNP1, revealing a marked difference in the transcription of numerous genes related to immunometabolism under lipotoxic conditions (**Figure S7a**). In fact, the RNA-seq profile of FFA-treated primary hepatocytes more closely resembled that of untreated cells than that of cells treated with FFA plus HNP1 (**Figure S7a**), underscoring the importance of HNP1 for perturbing the immune-metabolic pathways. GO analysis enriched top differential genes pertinent to lipid metabolism, energy homeostasis, inflammatory response, and neutrophil chemotaxis (**Figure S7b**). Network analysis further pinpointed enrichment to MASLD and inflammatory response, with the former closely coupled with mitochondrial processes (*e.g.,* mitochondrial protein-containing complex, mitochondrial envelope, voltage ion channel regulator activity and protein dimerization activity) (**Figure S7c**).

As an amphipathic molecule, cationic HNP1 may possess the ability to target the negatively charged mitochondrial envelope^52,53^. Indeed, FAM-labelled HNP1 could efficiently traverse BNL CL.2 cells to colocalize with mitochondria stained by MitoTracker Red (**Figure 3d**). Liver tissues, isolated from WT and *HNP1^TG^* mice after 12-week HFD feeding, were subjected to analysis of mitochondrial homeostasis using transmission electron microscopy (TEM). As shown in **Figure 3e**, hepatocytes from *HNP1^TG^* mice displayed an altered mitochondrial morphology and/or excessive damage as evidenced by disappeared cristae, reduced size and worsened fragmentation compared with WT controls (**Figure S8a**). These findings on HFD-fed *HNP1^TG^* mice were fully corroborated by results on clinical samples (*n = 6* in non-steatosis group; *n = 6* in steatosis group), where hepatic mitochondria from patients with hepatic steatosis were less uniform and more fragmented than those from patients without hepatic steatosis (**Figure 3f and Figure S8b**). Together, these findings strongly suggest that HNP1 directly interacts with hepatocytic mitochondria to induce mitochondrial dysfunction, a hallmark of hepatic steatosis.

### HNP1 targets the N-terminal helical domain of VDAC1 to promote its oligomerization leading to dysregulated lipid metabolism in hepatocytes

α-Defensins are capable of interacting with diverse molecular targets to exert their multiplicity of biological functions^46^, including with membrane-embedded host receptors to modulate innate immune responses^32,54^. To identify HNP1’s interacting molecule in mitochondria, we focused on our primary “suspect,” voltage-dependent anion channel 1 (VDAC1), the most abundant mitochondrial envelope protein essential for homeostasis via the maintenance of membrane potential and redox balance^40^. Our “suspicion” was reinforced by our transcriptomic analysis, which showed enrichment of the pathway associated with voltage-gated ion channel regulator activity (**Figure S7c**). Indeed, HNP1 was found to colocalize with the VDAC1 protein on liver tissues isolated from clinical patients (**Figure 4a**). To validate their direct interaction, VDAC1, a β-barrel protein with an N-terminal helical domain (**Figure 4b**), was expressed and purified (**Figure S9**), and subjected to a fluorescence polarization assay with FAM-labelled HNP1. As shown in **Figure 4c**, a VDAC1 dose-dependent increase in polarization values yielded, after a nonlinear regression analysis^30,55^, a K_d_ value of 83.4 nM, thereby confirming a potent binding affinity of HNP1 for VDAC1.

**Figure 4.**
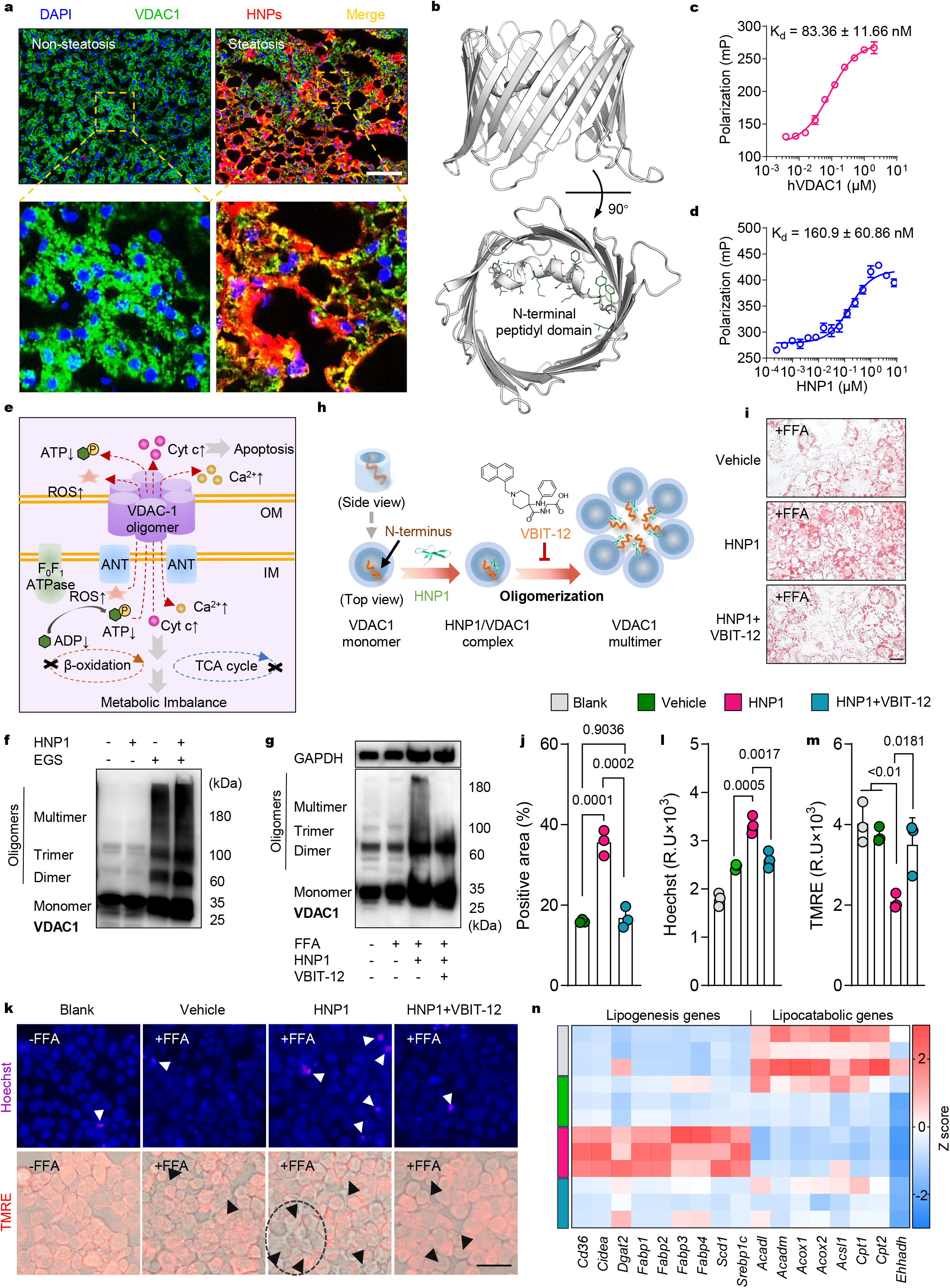
HNP1 targets the N-terminal helical domain of VDAC1 to promote its oligomerization leading to dysregulated lipid metabolism in hepatocytes. **(a)** Immunofluorescent images of liver sections from clinical patients with or without hepatic steatosis. Blue, nucleus; green, VDAC1; red, HNP1. Scale bar: 100 μm. **(b)** Crystal structure of human VDAC1 (PDB: 6G6U). (c-d) Binding affinity of HNP1 to full-length human VDAC1 **(c)** and to the 26-amino acid VDAC1 N-terminal peptide **(d),** assessed by fluorescent polarization. Data are presented as mean ± SEM from three biological replicates. **(e)** Schematic diagram of the mechanistic and functional outcomes resulted from VDAC1 oligomerization. **(f)** HNP1-induced oligomerization of purified human VDAC1 was visualized by immunoblotting after treatment with the cross-linking reagent EGS to stabilize the oligomers during electrophoresis. **(g)** VDAC1 oligomerization in mouse primary hepatocytes in the presence or absence of 0.5 mM palmitic acid, 4 μM HNP1, or 20 μM VBIT-12 was visualized by immunoblotting. **(h)** Schematic diagram of the proposed model in which HNP1 promotes VDAC1 oligomerization by interacting with its N-terminal domain. **(i)** Representative images of ORO-stained hepatocytes treated with 0.5 mM palmitic acid in the presence or absence of 4 μM HNP1 and/or 20 μM VBIT-12. Scale bar, 100 μm. **(j)** Semi-quantitative analysis of lipid accumulation in hepatocytes based on the percentages of ORO-positive area (*n* = 3 biological replicates). Data are shown as mean ± SD. Statistical significance was analyzed by one-way analysis of variance with the Bonferroni correction for multiple comparisons. **(k)** Representative images of Hoechst (apoptosis)- and TMRE (mitochondrial potential)-stained hepatocytes without (blank) or with FFA treatment in the presence of vehicle as control, 4 μM HNP1, or 4 μM HNP1 plus 20 μM VBIT-12. White arrows, apoptotic hepatocytes. Black arrows, hepatocytes with membrane potential depolarization. Scale bar: 50 μm. (**l-m**) Semi-quantitative analysis of fluorescence intensities of Hochest (**l**) and TMRE (**m**) in hepatocytes (*n* = 3 biological replicates). Data are shown as mean ± SD. Statistical significance was analyzed by one-way analysis of variance with the Bonferroni correction for multiple comparisons. (**n**) Transcriptional levels of genes related to lipogenesis and lipid catabolism in hepatocytes with various treatments. Relative mRNA levels were calculated as the quantification relative to blank controls according to the 2^-ΔΔCT^ mathematical model. The final data were normalized by Z-score and displayed as Z-scores in the microarray heatmap. n = 3 biological replicates per group.

To pinpoint HNP1’s binding site on VDAC1, we fluorescently labelled the C-terminus of a chemically synthesized peptide of 26 amino acid residues spanning the N-terminal helical domain of VDAC1 (NTP) and quantified its interaction with HNP1 using the fluorescence polarization technique. As shown in **Figure 4d**, HNP1 bound to the NTP peptide with a K_d_ value of 161 nM, largely recapitulating the binding affinity of HNP1 for full-length VDAC1 (**Figure 4c**) and implicating the N-terminal helical domain of VDAC1 as the molecular moiety directly targeted by HNP1.

Importantly, VDAC1 is prone to oligomerize under oxidative stress conditions (such as in a lipotoxic environment) to form large multifunctional channels or pores in the mitochondrial outer membrane, leading to the transport between the mitochondria and cytosol of many different types of molecules such as Ca^2+^, mtDNA, cytochrome c (Cyt c), and reactive oxygen species (ROS) (**Figure 4e**). When dysregulated, these cellular events can disrupt mitochondrial homeostasis, resulting in, among many others, altered metabolic plasticity, impaired lipid oxidation, lipoapoptosis, and exacerbated insulin resistance in conditions like diabetes and MASLD/MASH^41–43^. Since the evolutionarily conserved N-terminal helical domain of VDAC1 functionally regulates VDAC1 multimerization through governing its mobility^41,56^, it is plausible that HNP1, given its known ability to dimerize, oligomerize, and multimerize upon target binding^46^, may function as a scaffold to promote VDAC1 oligomerization. We verified this by showing that HNP1 indeed boosted the formation of VDAC1 dimers, trimers and high-order multimers *in vitro* (**Figure 4f**). More importantly, VDAC1 oligomerization was markedly enhanced in primary mouse hepatocytes cultured in the presence of 0.5 mM FFA and 4 μM HNP1 compared to those without HNP1 treatment, which was reversed by a VDAC1 oligomerization inhibitor, VBIT-12 (**Figure 4g**-**h**)^40^.

To ascertain HNP1-driven multimerization of the NTP peptide at the molecular level, we employed TEM technique to visualize the solution ultrastructure of NTP, HNP1, and their equimolar mixture, each at a concentration of 0.5 μM (**Figure S10a**). The NTP peptide and HNP1 alone formed well-dispersed miniature and moderately sized nanoparticles in aqueous solution, respectively, whereas their co-incubation gave rise to higher-order oligomers with enlarged volumes (**Figure S10a**). Additionally, we measured the hydrodynamic diameter (*d*_h_) distribution of each peptide and their equimolar mixture at concentrations of 0.1, 0.5, 2.5, or 12.5 μM using dynamic light scattering. Both NTP and HNP1 displayed a dose-dependent increase in size distribution (**Figure S10b**), biophysically corroborating the role of NTP domain in driving VDAC1 oligomerization and the intrinsic self-association tendency of α-defensins^41,46,56^. In agreement with the TEM findings, co-incubation of HNP1 and NTP at all four concentrations tested resulted in markedly enlarged *d*_h_ distributions (**Figure S10b**), indicating a productive interaction and supramolecular assembly between the two peptides in solution.

As expected, VBIT-12 effectively reduced HNP1-potentiated hepatic steatosis *in vitro* (**Figure 4i-j**), reinforcing the mechanistic importance of VDAC1 multimerization for HNP1-induced mitochondrial dysregulation of lipid metabolism. Notably, HNP1 also promoted considerable hepatocytic apoptosis and mitochondrial membrane potential depolarization under lipid-rich conditions *in vitro*, which was abolished by co-treatment with VBIT-12 (**Figure 4k-m**). In addition, RT-qPCR quantification of several genes involved in lipogenesis and lipid catabolism in primary mouse hepatocytes indicated that genes related to lipid catabolism were significantly downregulated at the transcriptional level in FFA-treated hepatocytes compared to cells without FFA treatment (**Figure 4n**). In contrast, HNP1 markedly upregulated lipogenesis-related genes in FFA-treated hepatocytes (**Figure 4n**). However, when VBIT-12 was added to FFA-treated cells in the presence of HNP1, the transcriptional profiles were restored to the levels comparable to those observed with FFA alone (**Figure 4n**). Collectively, these findings demonstrate that HNP1 predominantly interacts with the N-terminal helical domain of VDAC1 to promote VDAC1 multimerization, resulting in mitochondrial dysfunction and impaired lipid metabolism in hepatocytes.

### Structural basis for HNP1-VDAC1 interactions

To probe the structural determinants by which HNP1 interacts with the N-terminal helical domain of VDAC1, we expressed and purified an ^15^N-labelled NTP peptide (**Figure S11**) for analysis by 2D ^1^H-^15^N heteronuclear single quantum correlation (HSQC) NMR spectroscopy in the absence or presence of wild-type HNP1. As shown in **Figure 5a**, NTP displayed a characteristic α-helical conformation where its well-dispersed resonance peaks distributed between 7.3 and 9.0 ppm in the proton dimension. The addition of HNP1 to NTP at a molar ratio of 0.2:1 in 10 mM phosphate buffer (PB) alone significantly attenuated and/or shifted several resonance peaks at a consistent contour level (**Figure 5a**, black circles). In the presence of 40% TFE, the resonance peaks in the NTP spectra became more dispersed than those observed in 10 mM PB alone, indicating an expected enhancement in α-helicity. Although HNP1-induced signal attenuation was not observed under this condition, pronounced chemical shifts were detected for multiple resonance peaks (**Figure 5a**, black outlines). These findings collectively indicate a productive solution interaction between NTP and HNP1. However, this interaction was abolished by 0.2% sodium dodecyl sulfate (SDS), as demonstrated by the complete superimposition of the NMR spectra of NTP in the absence and presence of HNP1 (**Figure 5a**).

**Figure 5.**
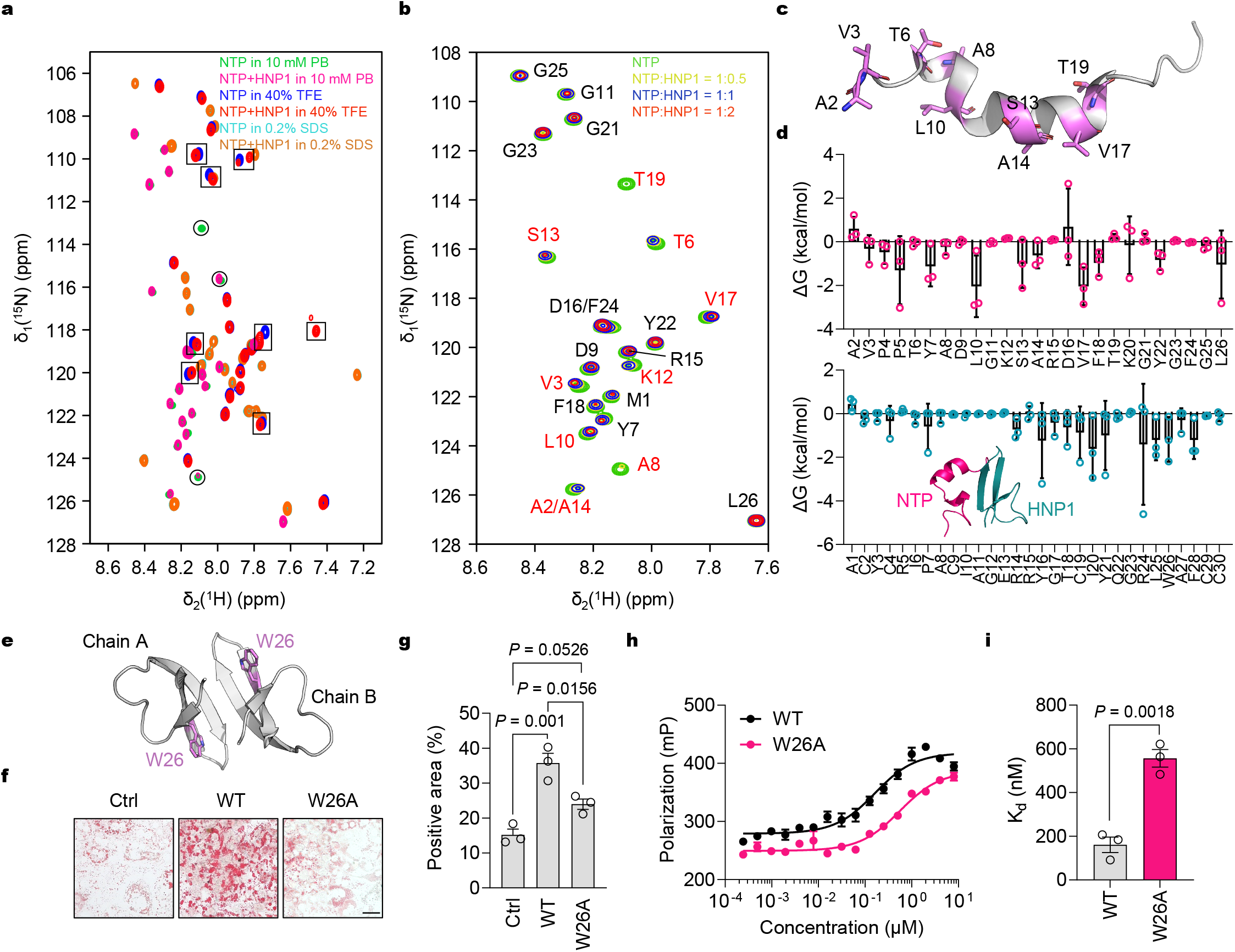
Structural basis for HNP1 binding to the VDAC1 N-terminal oligomerization domain. (**a**) 2D [^15^N, ^1^H] HSQC spectra of 400 μM ^15^N-labeled NTP in the absence and presence of 80 μM HNP1 under different buffer conditions. (**b**) 2D [^15^N, ^1^H] HSQC spectra of 200 μM ^15^N-labeled NTP superimposed with spectra of NTP titrated with 0.5, 1, or 2 equivalents of HNP1. (**c**) NTP structure extracted from crystal structure of human VDAC1 (PDB: 6G6U) with critical residues shown as stick. (**d**) Contributions of residues in NTP (top) and HNP1 (bottom) to MM/GBSA binding free energy. Data are presented as mean ± SD. (**e**) Crystal structure of HNP1 dimer (PDB: 3GNY). (**f-g**) Representative images of ORO-stained primary hepatocytes treated with 0.5 mM palmitic acid in the presence or absence of 4 μM HNP1 or W26A-HNP1 (**f**), and semi-quantitative analysis (**g**) of lipid accumulation in hepatocytes based on the percentages of ORO-positive area (*n* = 3 biological replicates). Scale bar, 100 μm. Data are shown as mean ± SEM. Statistical significance was analyzed by one-way analysis of variance with the Bonferroni correction for multiple comparisons. (**h-i**) Binding affinity (**h**) and K_d_ values (**i**) of the VDAC1 N-terminal peptide to HNP1 and its W26A mutant, assessed by fluorescent polarization. Data are shown as mean ± SEM. Statistical significance was analyzed by one-way analysis of variance with the Bonferroni correction for multiple comparisons.

Notably, when 2D ^1^H-^15^N HSQC spectra were collected for 200 μM NTP in 10 mM PB, either alone or after the addition of 0.5, 1, or 2 equivalents of HNP1 (**Figure 5b**), HNP1 dose-dependently reduced the signal intensity of most resonance peaks in the NTP spectra, especially those of A2, V3, T6, A8, L10, K12, S13, A14, V17, and T19 (**Figure 5b**), suggesting HNP1-dependent oligomerization of NTP. Consequently, the resonance peaks of A2, V3, T6, A14, V17 also shifted significantly (**Figure 5b**). These NMR data enabled experimental identification of critical residues in the VDAC1 N-terminal domain required for HNP1 engagement (**Figure 5c**).

For additional structural validation, HNP1 and NTP molecules were randomly placed and solvated in a rectangular box for subsequent 200-ns molecular dynamics (MD) simulations. Per-residue contributions for NTP to HNP1 binding were calculated using the MM/GBSA method based on MD trajectories (**Figure 5d**), which largely corroborated the NMR results. Of note, the C-terminal residues of HNP1 contributed the most to the binding free energy for its interactions with NTP (**Figure 5d**), in agreement with the fact that the functional determinants of HNP1 are heavily clustered in its C-terminal region with Trp26 being the most important^57^. For verification, we quantified the binding affinity of the NTP peptide for W26A-HNP1 using the fluorescence polarization assay and subjected the defensin analog to the *in vitro* lipotoxicity assay, where primary mouse hepatocytes were cultured under lipotoxic conditions in the presence of W26A-HNP1, followed by ORO staining and semi-quantitative analysis. As shown in **Figure 5e-i**, the W26A mutation not only significantly weakened HNP1 binding to NTP, but also substantially reduced defensin-stimulated lipid deposition in hepatocytes, thus confirming the functional importance of Trp26 for HNP1 as reported^57^.

### Oral administration of a VBIT-12 prodrug prevents the development of MASLD in *HNP1^TG^* mice fed with HFD

Encouraged by the *in vitro* activity of VBIT-12 in reducing lipid deposition, we set out to test its efficacy against MASLD *in vivo* as a proof of concept of leveraging the “HNP1-VDAC1 oligomerization” axis to attain a new therapeutic paradigm. To facilitate oral administration, we also synthesized a VBIT-12 prodrug (VBIT-12^pro^) via esterification of the free carboxyl group in VBIT-12 (**Figure S12**). This modification to VBIT-12 was necessary as it significantly increased lipophilicity, enabling the prodrug to passively diffuse across intestinal membranes for efficient absorption by the liver where endogenous carboxylesterases would ultimately cleave the ester bond to release active VBIT-12. As shown in **Figure S13**, VBIT-12 and VBIT-12^pro^ in particular showed negligible cytotoxicity against THLE-2 and BNL CL.2 cell lines at the highest concentration of 100 μM tested.

*HNP1^TG^* mice that had undergone six weeks of HFD feeding were then subjected to a daily oral administration (p.o.) for another 6 weeks of 20 mg/kg VBIT-12^pro^ or a vehicle control under continued HFD feeding, alongside mice receiving a daily intraperitoneal (i.p.) injection of 20 mg/kg VBIT-12 or a vehicle control in an otherwise identical head-to-head comparison (**Figure 6a**). As expected, all groups of mice showed similar velocity of ponderal growth during the initial 6-week HFD feeding period (**Figure 6b** and **Figure S14a-b** and **S15a**). However, their growth trajectories started to diverge upon initiation of treatment, with VBIT-12 or VBIT-12^pro^ treatment groups showing markedly slowed weight gain compared to vehicle controls during the subsequent 6 weeks of HFD feeding (**Figure S14c and S15b**). In fact, the body weights of *HNP1^TG^* mice treated with VBIT-12^pro^ were approximately 8.65% and 13.8% lower, respectively, than those of vehicle controls after 8 and 12 weeks of HFD feeding (**Figure 6c**).

**Figure 6.**
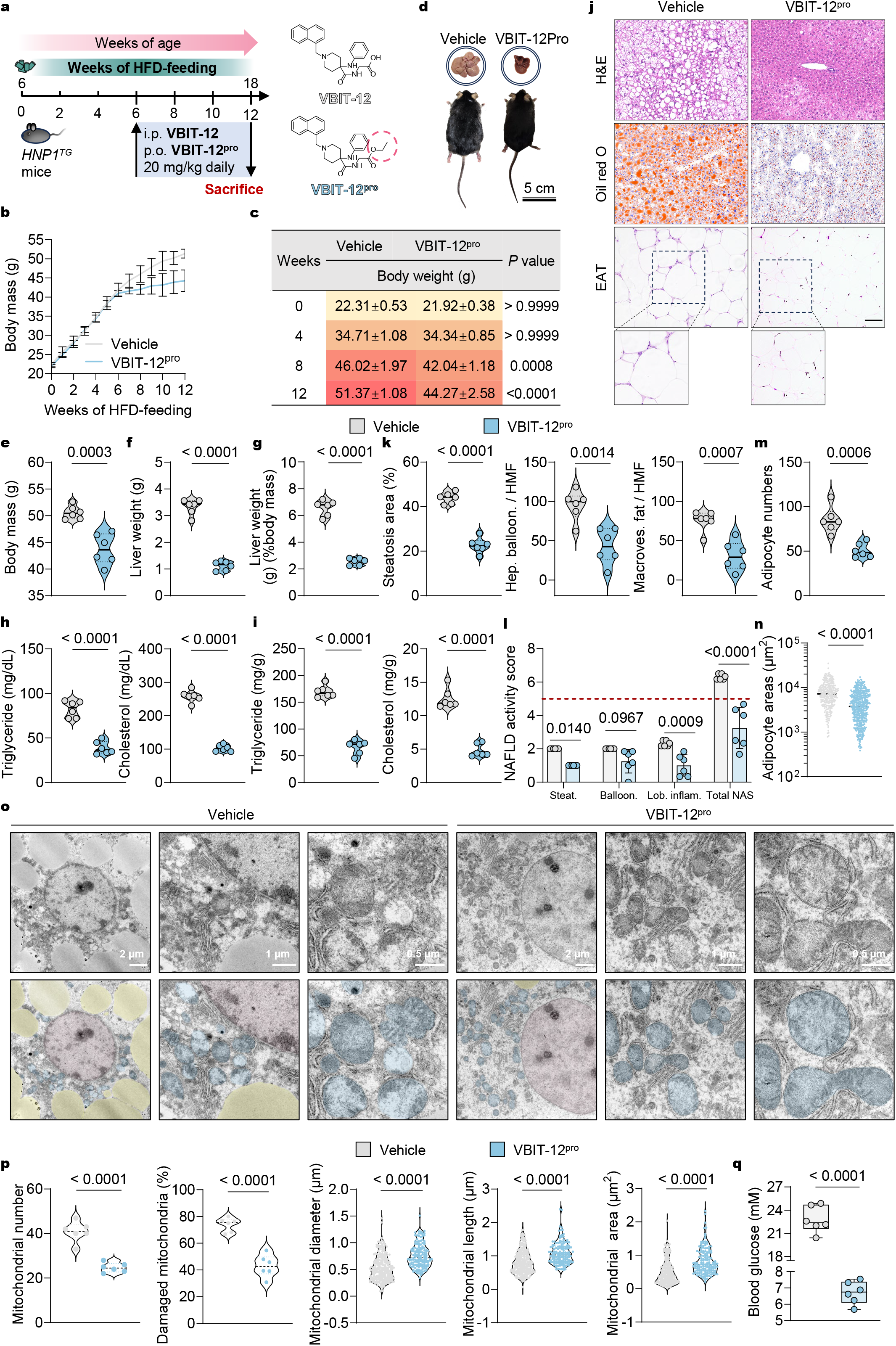
Relieved MASLD in *HNP1^TG^* mice by oral administration of VBIT-12 prodrug. (**a**) Experimental design for evaluating the *in vivo* efficacy of VBIT-12 and VBIT-12^pro^ against MASLD in *HNP1^TG^* mice. Six-week-old male *HNP1^TG^* C57BL/6J mice, weighed 21-23 g, were fed an HFD for 6 weeks without medication, followed by another 6 weeks of HFD feeding with daily intraperitoneal injection (i.p.) of VBIT-12 or oral administration (p.o.) of VBIT-12^pro^ at a dose of 20 mg/kg, after which they were sacrificed and tissue samples collected for in-depth analysis. (**b**) Weekly body weights of HFD-fed *HNP1^TG^* mice treated with vehicle or VBIT-12^pro^ (*n* = 6 mice per group). (**c**) Summary of body weights and statistical comparisons between vehicle- and VBIT-12^pro^-treated *HNP1^TG^* mice at 0, 4, 8, and 12 weeks. (**d**) Representative photographs of *HNP1^TG^* mice treated with vehicle or VBIT-12^pro^ (bottom) and their livers (top) following 12 weeks of HFD feeding. (**e-g**) Body weights (**e**), liver weights (**f**), and liver weight/body weight ratios (**g**) of vehicle- and VBIT-12^pro^-treated *HNP1^TG^* mice after 12 weeks of HFD feeding (*n* = 6 mice per group). (**h-i**) Levels of triglyceride and cholesterol in serum (**h**) and livers (**i**) from vehicle- and VBIT-12^pro^-treated *HNP1^TG^* mice after 12 weeks of HFD feeding (*n* = 6 mice per group). (**j**) Representative images of H&E- and ORO-stained mouse liver sections, and H&E-stained mouse EAT sections. Scale bar: 100 μm. (**k**) Quantification of ORO-positive area percentage (left), ballooned hepatocytes (middle), and macrovesicular fat-laden hepatocytes (right) per HMF using ImageJ, based on three fields per section (*n* = 6 mice per group). (**l**) NAFLD activity score (NAS) of livers from vehicle- or VBIT-12^pro^-treated *HNP1^TG^* mice after 12 weeks of HFD feeding (*n* = 6 mice per group). (**m**) Adipocyte number count in EAT sections using ImageJ, based on three fields per section (*n* = 6 mice per group). (**n**) Quantification of adipocyte area in H&E-stained EAT sections from vehicle-and VBIT-12^pro^-treated *HNP1^TG^* mice (305 and 510 adipocytes measured, respectively). (**o-p**) Representative TEM images (**o**) of hepatocellular mitochondria and lipid droplets in livers of HFD-fed *HNP1^TG^* mice treated with vehicle or VBIT-12^pro^ and quantification of mitochondrial parameters based on their TEM images (**p**). (**q**) Levels of fasting blood glucose in serum from *HNP1^TG^* mice that were fed an HFD for 12 weeks and received daily treatment with vehicle or VBIT-12^pro^ for the last 6 weeks. All data are shown as mean ± SD. Statistical significance was analyzed by an unpaired, 2-tailed Student’s *t* test.

VBIT-12 or VBIT-12^pro^ treatment not only retarded the runaway growth of *HNP1^TG^* mice fed with HFD, but also reduced their liver volume to normal dimensions (**Figure 6d and S14d**). Upon conclusion of the six-week treatment, mice were sacrificed after overnight fasting, followed by collection of livers and EAT pads for further analysis. The body weights of *HNP1^TG^* mice medicated with VBIT-12^pro^ decreased by approximately 14% (**Figure 6e**), and their liver weights were only one-third those of the vehicle controls (**Figure 6f-g**), whereas the EAT pads from VBIT-12^pro^-treated mice increased by about 46% (**Figure S15c-d**). Furthermore, both triglyceride and cholesterol levels in serum and liver dropped because of VBIT-12^pro^ treatment (**Figure 6h-i**), which correlated with attenuated liver injury, as evidenced by reduced ALT and AST levels (**Figure S15e-f**).

Notably, oral gavage of VBIT-12^pro^ significantly reduced ballooned and macrovesicular hepatocytes and alleviated hepatic steatosis, as shown by H&E and ORO staining (**Figure 6j-k**). All mice in the vehicle control groups progressed to the MASH stage (NAS > 5), whereas VBIT-12^pro^ effectively reversed disease progression from MASH to MASL (**Figure 6l**). In addition, we observed smaller adipocytes with fewer CLS in the EAT pads of VBIT-12^pro^-treated mice compared to vehicle controls (**Figure 6j, m and n**). To assess mitochondrial physiology, we examined the ultrastructure of mitochondria in mouse livers using TEM imaging. In the liver tissues from vehicle-treated controls, we observed numerous circular lipid droplets (yellow) surrounding the nucleus (red), along with mitochondrial fragmentation and invaginated cristae (blue) (**Figure 6o**). In contrast, the livers of VBIT-12^pro^-treated mice displayed apparently reduced lipid droplet infiltration and intact mitochondria with distinct double membranes and well-preserved cristae (**Figure 6o**). These observations were further reinforced by quantitative analysis of mitochondrial number and damage severity (**Figure 6p**). Not surprisingly, blood glucose levels in *HNP1^TG^* mice decreased to normal range upon VBIT-12^pro^ treatment (**Figure 6q**). Although intraperitoneal injection of VBIT-12 achieved similar therapeutic efficacy against MASLD (**Figure S14e-u**), the orally available prodrug VBIT-12^pro^ likely offers a greater translational potential as a new class of therapeutic agents for the treatment of MASLD and possibly other metabolic disorders arising from premature multimerization, forced or otherwise, of VDAC1 in hepatocytes.

## DISCUSSION

Persistent infiltration of neutrophils recruited by Kupffer cells to the site of hepatic inflammation under conditions of lipid overload contributes to disordered lipid metabolism and hepatocyte death seen in MASLD^15,58^, and, yet the precise molecular mechanisms of action of neutrophils remain only partially understood. Neutrophil α-defensins as important host protective factors are stored in azurophilic granules at a local concentration of 10-50 mg/mL and released to phagosomes to kill engulfed microbes^28,29^. Despite that hundreds of milligrams of α-defensins are produced daily in an adult human being by 5× 10^10^-10×10^10^ neutrophils with a half-life of 6-8 h^59^, only a tiny fraction of these peptides is found in one’s blood plasma and tissue fluids^60,61^. Where the bulk of these defensins are disseminated from dying neutrophils largely remains unknown, as is the case with their potential physiological or pathological functions at likely sub-bactericidal or sub-cytotoxic concentrations^27,62^. We have discovered that human neutrophil α-defensins HNPs, significantly elevated in the liver of clinical patients with hepatic steatosis, efficiently traverse hepatocytes to induce VDAC1 multimerization, thereby promoting hepatocytic mitochondrial dysfunction, lipid deposition and, ultimately, MASLD development and progression – a process that nevertheless can be reversed by inhibitors of VDAC1 multimerization.

Our unprecedented findings are significant at multiple levels. First, human neutrophil α-defensins best known for their antimicrobial and immunomodulatory activities in innate immunity are now endowed with a new pathological function in lipid metabolism and MASLD, reinforcing the tenet that these host defense peptides can act as a “double-edged sword” in health and disease^33^. Second, although hepatocytic mitochondria, a central hub for fatty acid metabolism via β-oxidation, is known to play a critical role in MASLD^63^, our studies unveil an unexpected mechanism of MASLD development and progression by which neutrophil α-defensins in the liver induce mitochondria dysfunction to trigger a cascade of biological events leading to exacerbated hepatic steatosis and worsened MASLD. Third, central to this newly discovered molecular mechanism that drives MASLD pathogenesis is the mitochondrial membrane protein VDAC1 multimerized by neutrophil α-defensins, which conceptualizes a highly attractive therapeutic paradigm for MASLD. Fourth, our studies indeed cultivate the design of an orally administered prodrug, based on the known inhibitor of VDAC1 multimerization VBIT-12^40^, with superior *in vitro* and *in vivo* pharmacological efficacy in treating MASLD, which, to our best knowledge, is likely a first-in-class drug candidate of this sort with significant translational potential.

Human neutrophil α-defensins can play other pathological roles under certain *in vitro* and/or *in vivo* biological settings. HNP1, for example, can form heteromers with the platelet-derived chemokine CCL5 to augment monocyte recruitment and activation in acute and chronic inflammation, exacerbating myocardial infarction in mice overexpressing both proteins^64^. Neutrophil α-defensins have also been shown to cause lung injury by disrupting the capillary-epithelial barrier in HNP1/HNP2-transgenic mice^65^ and to induce sepsis-related organ damage and mortality in HNP1/HNP3-transgenic mice through endothelial cell pyroptosis^54^. A more recent report shows that sublethal concentrations of HNPs released by neutrophils at the site of *A. baumannii* infection in the respiratory tract directly interact with the outer membrane protein OmpA to promote bacterial biofilm formation and antibiotic tolerance^30^. Further, these neutrophil α-defensins can also enhance *Shigella* adhesion to and invasion of host cells^66^. In fact, a structurally homologous enteric antimicrobial peptide produced by Paneth cells in the small intestine^67^, human α-defensin 5 (HD5), enhances *Shigella* infection through two independent mechanisms, (1) HD5-OmpA-mediated bacterial adhesion and invasion^31^ and (2) HD5-P2Y11-activated cytoskeleton rearrangement and extension of filopodia in colonic epithelial cells for bacterial capturing^32^. Importantly, these pathological functions of human α-defensins appear pertinent, as most of their antimicrobial and immunomodulatory activities, to their ability to dimerize, oligomerize and multimerize in solution, which is intensified by the presence of their promiscuous molecular targets of both microbial and host origins^46,47,68^.

Of note, LL-37, the sole member in humans of the cathelicidin family of antimicrobial peptides expressed by neutrophils, macrophages and epithelial cells^69–71^, has also been implicated as a culprit in autoimmune disorders such as psoriasis, systemic lupus erythematosus and rheumatoid arthritis due to its potent immunomodulatory properties^72,73^. More recently, Nakamura et al. reported that LL37 promotes LDL uptake in macrophages through multiple receptors, increasing risks of atherosclerosis in patients with chronic inflammatory disorders whose cathelicidin levels are elevated^74^. However, while circulating LL37 levels in humans are associated with obesity, overexpression of LL37 in mice reduces hepatic steatosis and suppresses lipid accumulation in adipocytes and hepatocytes by inhibition of the CD36 fat receptor^75^, thus contrasting our findings with HNPs.

It is plausible that aberrant VDAC1 multimerization impairs mitochondrial function and exacerbates hepatocytic lipid deposition through multiple interconnected mechanisms. First, it may compromise fatty acid β-oxidation, thereby reducing the liver’s capacity to dispose of excess energy derived from processed foods^6,43^. Second, since VDAC1 dimers function as scramblase-type lipid transporters that catalyze lipid translocation across the outer membrane to support mitochondrial function^76^, it may perturb mitochondrial membrane homeostasis^77^. Third, it may increase outer membrane permeability, facilitating mtDNA release and subsequent NLRP3 inflammasome activation, which in turn exacerbates lipid deposition in hepatocytes^44,78–81^. Despite the prior identification of Pemt and BRD4 as potential upstream regulators of VDAC1 association^44,45^, the direct molecular drivers of VDAC1 multimerization, in addition to HNPs identified in this study, remain to be fully elucidated, particularly under pathological conditions such as MASLD.

Of note, under physiological settings, mitochondria in hepatocytes undergo a dynamic fusion-fission cycle critical for sustaining mitochondrial homeostasis and metabolic coordination^82–84^. Hence, MASLD development is invariably accompanied by disruption of mitochondrial morphology and integrity. Indeed, we observed significantly altered hepatocyte mitochondrial ultrastructure with cristae disappearance and other defects in liver tissues from HFD-fed *HNP1^TG^* mice, but not in WT mice under the same feeding conditions. Moreover, compared to WT mice, *HNP1^TG^* mice exhibited exacerbated hepatic steatosis along with reduced EAT pads after 12 weeks of HFD feeding, suggesting a potential role of adipose tissue remodeling. This may result from worsened insulin resistance and inflammation in the white adipose tissue of *HNP1^TG^* mice, which redirect ingested glucose and fatty acids to the liver, thereby imposing an additional metabolic burden on hepatocytes^6^.

That human neutrophil α-defensins promote MASLD development and progression by inducing mitochondrial dysfunction establishes an unexpected but definitive link between innate immunity and lipid metabolism, underscoring the largely underappreciated functional pleiotropy of these host factors beyond immune responses to microbial infections. Future studies are warranted to (1) delineate at the atomic level the molecular basis for HNPs-induced VDAC1 multimerization using structural tools such as NMR spectroscopy, X-ray crystallography, and cryo-electron microscopy, (2) determine whether HNPs can further drive the progression of hepatic steatosis to cirrhosis and HCC using long-term HFD-fed *HNP1^TG^* mice, and (3) decipher whether HNPs, other defensins or innate effectors may contribute more broadly to the metabolic syndrome or a disease setting through regulating cell metabolism or mitochondrial dynamics. Finally, since mitochondrial VDAC1 is highly expressed in metabolically active tissues other than liver, such as malignant tumors, brain, and muscle^40,85,86^, HNPs released from neutrophils in response to infection, inflammation and sterile injury may play previously unrecognized pathological roles that have yet to be fully elucidated. In this regard, inhibitors of VDAC1 multimerization such as VBIT-12^pro^ may also prove useful as a new class of therapeutic agents for the treatment of cancer, neurodegenerative, and cardiovascular diseases.

## DATA AVAILABILITY

All other data are available in the main text or the supplementary materials. Resources, reagents and materials generated in this study are available from the Lead contact, W. L., upon request.

## CODE AVAILABILITY

This paper does not report original code.

## ACKNOWLEDGEMENTS

We thank Dr. Lingyun Yang in NMR platform of iHuman Institute, ShanghaiTech University, for his excellent technical assistance on 2D NMR experiments. This work was supported by grants from the National Key Research and Development Program of China (2024YFA1803102 to W. L.), the National Natural Science Foundation of China (82400683 to J. Z., 82030062 to W. L., and 82604424 to G. L.), the China Postdoctoral Science Foundation (2023M740661 to J. Z and 2026M791501 to G. L.), the China National Postdoctoral Program for Innovative Talents (BX2026366 to G. L.), and the Shanghai Municipal Science and Technology Major Project (ZD2021CY001 to W. L.).

## AUTHOR CONTRIBUTIONS

W. L., G. L., and J. Z. conceived, supervised, and/or designed the research. X. L. collected all clinical samples involved in this study. J. Z., G. L., G. W., and J. R. performed all experiments and characterizations. G. L. and J. Z. wrote the original draft of this manuscript. W. L. extensively revised this manuscript.

