## Supplementary material for "A Neutrophil Defensin–VDAC1 Axis Drives Steatotic Liver Disease": Methods and Supplemental figures 1-15

**Table of contents:**

Methods

References

Figure S1-S15

### METHODS

**Human liver tissues.** Clinical liver specimens were sourced from the biological sample bank of the First Affiliated Hospital of the Air Force Medical University, with ethical approval granted by the hospital's Medical Ethics Committee (approval number: KY20232280-X-1). Based on oil red O (ORO) staining quantification, specimens were classified according to international guidelines into a non-NAFLD group (steatosis area <5%) and a NAFLD group, the latter further subdivided into mild (5%–33%) and severe (>33%) categories<sup>1,2</sup>.

**Immunohistochemistry and quantification of HNP1.** Formalin-fixed, paraffin-embedded liver sections from human and mouse were used to assess the presence and relative content of HNP1. Sections were incubated overnight at 4 °C with an anti-HNP1 primary antibody (1:2000, HUABIO, EM1701-52). After washing, sections were incubated with a horseradish peroxidase-conjugated anti-mouse secondary antibody for 30 min at room temperature, followed by hematoxylin counterstaining. Images were acquired using a BZ-X810 fluorescence microscope (KEYENCE), with six high magnification fields captured per section. The HNP1-positive area fraction in human livers was quantified with the IHC Profiler plugin in ImageJ. To normalize hepatic HNP1 content across specimens, the sample with the lowest positive area percentage was designated as the baseline (relative value

= 1), and fold changes for all other samples were calculated relative to this baseline. A semi-quantitative scoring system was applied to grade HNP1 expression: each section received a composite score calculated as the product of staining intensity (0 = none, 1 = weak, 2 = moderate, 3 = strong) and the proportion of positive cells (1 = 0–25%, 2 = 25–50%, 3 = 50–75%, 4 = 75–100%). Expression levels were classified as weak (score 0–3), moderate (score 3–6) or strong (score 6–12).

**Histopathological analysis.** Hematoxylin and eosin (H&E) staining was performed on 4 µm formalin-fixed, paraffin-embedded tissue sections. For assessment of hepatic lipid deposition, 10 µm frozen liver sections were stained with ORO. The NAFLD activity score (NAS) was calculated as the sum of individual scores for steatosis, ballooning, and lobular inflammation, according to published criteria<sup>1,2</sup>. Steatosis was scored as 0 (<5%), 1 (5–33%), 2 (33–66%), or 3 (>66%); ballooning as 0 (none), 1 (few balloon cells), or 2 (many/prominent balloon cells); and lobular inflammation as 0 (no foci), 1 (<2 foci), 2 (2–4 foci), or 3 (>4 foci). Quantitative morphometric analysis was performed using ImageJ on six high magnification fields per section. The following parameters were evaluated: (a) ballooned and macrovesicular hepatocytes on H&E-stained liver sections; (b) the percentage of lipid deposition area on ORO-stained liver sections; and (c) adipocyte number and cross-sectional area on H&E-stained epididymal adipose tissue (EAT) sections.

All histological images were captured with a BZ-X810 fluorescence microscope (KEYENCE).

**Fibrotic stage evaluation.** To assess hepatic fibrosis, formalin-fixed, paraffin-embedded liver sections were stained with Masson's trichrome and Sirius Red. Quantitative analysis was performed using ImageJ on six randomly selected high magnification fields per section. The fibrosis area fraction was measured on Masson's trichrome-stained sections, and collagen types were distinguished on Sirius Red-stained sections under polarized light. Images of Masson's trichrome staining were captured using a BZ-X810 fluorescence microscope (KEYENCE), whereas Sirius Red staining was visualized with a Polarizing Optical Ci upright microscope (Nikon, Japan).

**Animal models.** All animal procedures were reviewed and approved by the Animal Ethics Committee of the School of Basic Medicine, Fudan University (approval number: 20240229-039). To minimize variability due to hormonal fluctuations, only male mice were included. Animals were housed under specific-pathogen-free (SPF) conditions in standard cages.

To generate neutrophil-specific humanized mice expressing HNP1, the human *DEFA1* gene (NCBI Gene ID: 1667) was inserted into the H11 locus of C57BL/6J mice under the control of the neutrophil elastase (NE) promoter,

using CRISPR-Cas9 technology. This construct was produced by
GemPharmatech Co., Ltd. (Nanjing, China).

Approximately six-week-old wild-type (WT) and HNP1 transgenic (*HNP1<sup>TG</sup>*) mice, weighing 21–23 g, were fed either a control fat diet (CFD; 20 kcal% protein, 10 kcal% fat, 70 kcal% carbohydrates; Research Diets, D12450B,) or a high-fat diet (HFD; 20 kcal% protein, 60 kcal% fat, 20 kcal% carbohydrates; Research Diets, D12492) *ad libitum* for 12 weeks to establish control and NAFLD models, respectively. All mice were fasted overnight before tissue collection.

To evaluate the therapeutic efficacy of VBIT-12 and VBIT-12<sup>pro</sup> in the NAFLD model, six-week-old *HNP1<sup>TG</sup>* mice (body weight 21–23 g) were fed HFD for 12 weeks. After 6 weeks of HFD feeding, the mice were treated once daily for the subsequent 6 weeks with either VBIT-12 (20 mg/kg, i.p.) or VBIT-12<sup>pro</sup> (20 mg/kg, p.o.). At the end of the 6-week treatment period, the mice were fasted overnight before tissue harvesting.

**Mouse Genotype Identification.** Toe samples from *littermate* mice were collected and placed into centrifuge tubes containing 50 µL 50 mM NaOH, and heated 95 °C in a metal bath for 1 h. After cooling for 5 min, an equal volume of 100 mM Tris-HCl pH 7.4 was added and mixed. The resulting lysate (2 µL) was used as template for PCR amplification with 2 × Phanta Ultra Master Mix (P518, Vazyme, China) using the following primers:

H11-tF3: GGGCAGTCTGGTACTTCCAAGCT,

WPRE-q-tR1: TTGCGTCAGCAAACACAGTGCA;

WPRE-tF1: TCAATCCAGCGGACCTTCCTT,

H11-Tr3: ATATCCCCTTGTTCCCTTTCTGC;

H11-wt-tF1a: AGTCTTTCCCTTGCCTCTGCT,

H11-wt-tR1a: GGGTCTTCCACCTTTCTTCAG.

PCR products were separated by 2% agarose gel electrophoresis, and bands were visualized using a Tanon 1600 multifunctional gel image analysis system.

**Transcriptomics.** Transcriptome sequencing analysis was performed on clinical liver specimens, as well as liver tissues from 12-week HFD fed WT and *HNP1<sup>TG</sup>* mice, and mouse primary hepatocytes subjected to three treatments: untreated, 0.5 mM palmitic acid, and 0.5 mM palmitic acid with 4 μM HNP1. The transcriptomics service was provided by APEX-BIO. Following stringent quality control, gene expression profiling was conducted using the Illumina NovaSeq 6000 platform. Clean reads were aligned to the reference genome using HISAT2. Gene and transcript expression levels were quantified using StringTie. To screen for genes with significantly different expression levels in samples under different states. Differential expression analysis was performed with DESeq2 (for samples with biological replicates) or edgeR (for samples without replicates), using the read count data generated by StringTie. Gene Ontology (GO) functional enrichment and Kyoto Encyclopedia of Genes and Genomes (KEGG) pathway enrichment analyses were performed on differentially expressed gene (DEGs) sets using the clusterProfiler package. Pathway network analysis was conducted using Cytoscape following g:Profiler analysis (https://biit.cs.ut.ee/gprofiler/gost) of DEGs. Visualization of the other results was performed using online tools (https://www.bioinformatics.com.cn/aboutus, https://www.omicstudio.cn/tool) and GraphPad Prism 10.

**Metabolomics.** Liver tissues from 12-week HFD fed WT and *HNP1<sup>TG</sup>* mice were subjected to lipidomics analysis, which was performed by APExBIO. Briefly, samples were slowly thawed at 4 °C, and an appropriate amount of each sample was mixed with 200 µL of methanol and 10 µL of internal standard mixture. Subsequently, 800 µL of methyl tert-butyl ether (MTBE) was added and mixed. The mixture was ultrasonicated in an ice-cold water bath for 20 min and incubated at room temperature for 30 min. Then, 200 µL of mass spectrometry-grade water was added and vortexed. After centrifugation at 14000 rpm for 15 min at 4 °C, the upper organic phase was collected and evaporated to dryness under a nitrogen stream. The dried residue was reconstituted in 200 µL of 90% isopropanol/acetonitrile (v/v) solution, vortexed, and centrifuged again at 14000 rpm for 15 min at 4 °C. The supernatant was transferred for LC-MS/MS analysis.

Chromatographic separation was performed on an LC-30AD ultra-high-performance liquid chromatography (UHPLC) system (Shimadzu) using two types of columns. For the C18 column, the column temperature was maintained at 45 °C with a flow rate of 0.35 mL/min. Mobile phase A consisted of 70% acetonitrile, 30% water and 5 mM ammonium acetate; mobile phase B was isopropanol. The gradient elution program was as follows: 0–5.0 min, B increased linearly from 20% to 60%; 5.0–13.0 min, B increased linearly from 60% to 100%; 13.1–17.0 min, B was maintained at 20%. For the amino column, the column temperature was 40 °C with a flow rate of 0.4 mL/min. Mobile phase A consisted of 2 mM ammonium acetate in 50% methanol/50% acetonitrile; mobile phase B consisted of 2 mM ammonium acetate in 50% acetonitrile/50% water. The gradient program was: 0–3.0 min, B at 3%; 3.0–13.0 min, B increased linearly from 3% to 100%; 13.0–17.0 min, B at 100%; 17.1–22.0 min, B at 3%. Throughout the analysis, samples were maintained at 10 °C in the autosampler and analyzed in random order to minimize signal drift.

Mass spectrometry detection was performed using an AB 6500+ QTRAP mass spectrometer (AB SCIEX) equipped with an electrospray ionization (ESI)

source, operating in both positive and negative ion modes. The ESI source parameters were set as follows: source temperature, 400 °C; ion source gas 1 (GS1), 50; ion source gas 2 (GS2), 55; curtain gas (CUR), 35; ion spray voltage (IS), +3000 V (positive mode) or -2500 V (negative mode). Data acquisition was performed in multiple reaction monitoring (MRM) mode.

For the two-group comparison, differential metabolites were identified using orthogonal partial least-squares discriminant analysis (OPLS-DA) to obtain variable importance in projection (VIP) values, combined with unpaired, 2-tailed Student's *t* test to obtain *P* values. Fold change (FC) was calculated to determine the direction of regulation: metabolites with FC > 1 were considered upregulated, whereas those with FC < 1 were considered downregulated. A metabolite was considered significantly different if it simultaneously met the criteria of VIP > 1 and *P* < 0.05. Visualization of the results was performed using GraphPad Prism 10.

**Biochemical analyses.** Blood samples were allowed to clot at room temperature for 2 h and then centrifuged at 3,000 rpm for 10 min at 4 °C. Serum supernatants were collected and stored at -80 °C until analysis. Serum concentrations of alanine aminotransferase (ALT) and aspartate aminotransferase (AST) were determined using commercial kits (Nanjing Jiancheng Bioengineering Institute, ALT: C009-2-1; AST: C010-2-1;). Hepatic triglyceride and cholesterol contents were quantified using a Triglyceride Colorimetric Assay Kit (A111-1-1) and a Cholesterol Quantification Kit (A110-1-1), respectively (both from Nanjing Jiancheng), following the manufacturer's protocols. Blood glucose was measured with a glucose oxidase-based assay kit (Nanjing Jiancheng, A154-1-1).

**Glucose and insulin tolerance tests.** Glucose and insulin tolerance tests were performed on separate cohorts of WT and *HNP1<sup>TG</sup>* mice after 12 weeks of high-fat diet feeding. For GTT, mice were fasted overnight, weighed, and injected intraperitoneally with glucose (1.5 g/kg; Sinopharm Chemical Reagent, 63005518). For ITT, mice were fasted for 6 h, weighed, and injected intraperitoneally with recombinant human insulin (1 U/kg; TargetMol, T8221). In both assays, blood glucose levels were measured in tail vein blood using a glucometer (Cofoe, A03) at 0, 15, 30, 45, 60, 90 and 120 min after injection. Mice were euthanized upon completion of each test.

**Cell lines.** Mouse normal hepatocyte BNL CL.2 and human normal hepatocyte THLE-2 cell lines were purchased from the Type Culture Collection of the Chinese Academy of Sciences (Shanghai, China). Cells were maintained in Dulbecco's modified Eagle's medium (DMEM, Gibco, C11995500BT) supplemented with 10% fetal bovine serum (FBS, Gibco, 10099-141) and 1% penicillin–streptomycin (P/S, Gibco, 15140-122) at 37 °C in an incubator containing 5% CO<sub>2</sub>.

**Isolation and culture of primary hepatocytes.** Primary hepatocytes were isolated from mice following established protocols with modifications<sup>3,4</sup>. Briefly, mice were anesthetized by intraperitoneal injection of avertin (20 µL/g; Nanjing Aibei Biotechnology, M2910). The liver was perfused via the portal vein, first with perfusion buffer A (50 mL Krebs buffer containing 0.36 g/L KCl, 0.16 g/L KH<sub>2</sub>PO<sub>4</sub>, 0.3 g/L MgSO<sub>4</sub>, 7 g/L NaCl, 2 g/L NaHCO<sub>3</sub>, 3.6 g/L glucose, 1.2 g/L Hepes, and 0.1 mL of 50 mM EGTA), and then with perfusion buffer B (30 mL Krebs buffer containing 41.2 µL of 2 M CaCl<sub>2</sub> and 15 mg collagenase type I (Yeasen, 40507ES60)). The perfused liver was excised, minced, and filtered through a 200-µm mesh. Hepatocytes were washed with washing buffer (0.1 g/L DNase I (Sinopharm, 64002860;), 0.1 g/L MgSO<sub>4</sub>·7H<sub>2</sub>O and 0.1 g/L MgCl<sub>2</sub>·6H<sub>2</sub>O) and collected by centrifugation at 780 rpm for 2 min. Isolated hepatocytes were seeded onto plates pre-coated overnight with 50 µg/mL rat tail collagen I (Yeasen, 40125ES10) in 0.02 M acetic acid and cultured in complete DMEM medium (DMEM supplemented with 10% FBS and 1% penicillin–streptomycin) in a humidified incubator at 37 °C with 5% CO<sub>2</sub>. After 6 h, the medium was replaced with fresh complete DMEM medium for subsequent experiments.

**Cell treatment.** To establish an *in vitro* lipotoxicity model, BNL CL.2 mouse hepatocytes, primary mouse hepatocytes and THLE-2 human hepatocytes were treated with 0.5 mM palmitic acid (PA; Sigma, P0500) for 24 h. To evaluate the roles of HNP1 (or its variant) and VBIT-12 in hepatocyte steatosis, cells were incubated with 4 μM HNP1 or HNP1 variant, with or without 20 μM VBIT-12, under the lipotoxic conditions described above for 24 h.

**ORO staining of hepatocytes *in vitro*.** ORO staining of cultured hepatocytes was performed as described previously<sup>4</sup>. ORO stock solution was prepared by dissolving 0.5 g of ORO powder (Sangon Biotech, A600395) in 100 mL of isopropanol, with thorough mixing, and stored at 4 °C in the dark in a sealed container. Cells were fixed with 4% neutral buffered formaldehyde for 10 min, washed three times with PBS, and then incubated with ORO working solution (ORO stock: distilled deionized water = 3:2, filtered before use) for 20 min at room temperature. After staining, cells were rinsed with isopropanol solution (isopropanol: deionized water = 3:2) and maintained in PBS for image acquisition, and images were captured using a BZ-X810 fluorescence microscope (KEYENCE).

**Immunofluorescence.** Frozen human liver sections were fixed with 4% paraformaldehyde for 30 s, permeabilized with 0.25% Triton X-100 in PBS for 20 min at room temperature, and blocked with 3% BSA in PBS for 1 h. Sections were then incubated overnight at 4 °C with primary antibodies against human HNP1 (mouse monoclonal, 1:500, Proteintech, 67156-1-Ig) and human VDAC1/porin (rabbit monoclonal, 1:1,000, Abcam, ab154856). After washing, sections were incubated with appropriate fluorophore-conjugated secondary antibodies for 1 h at room temperature. Nuclei were counterstained with DAPI. Images were visualized by a fluorescence scanner KF-FL-020 (KFbio).

**Transmission electron microscopy (TEM) observation of mitochondrial ultrastructure.** For mitochondrial ultrastructure analysis, liver tissues were dissected and fixed in 2.5% glutaraldehyde overnight at 4 °C, post-fixed in 1% osmium tetroxide for 80 min at 4 °C, dehydrated through a graded ethanol series, and embedded in epoxy resin. Ultrathin sections (100 nm) were cut using an ultra-microtome, mounted on copper grids, and stained with 4% uranyl acetate and lead citrate. Images were acquired using a transmission electron microscope.

**Quantification of mitochondrial ultrastructural parameters.** Mitochondrial morphology was assessed on TEM images using ImageJ. The following parameters were measured for each mitochondrion: length, diameter, cross-sectional area, and number per field. For each sample, all mitochondria were counted and measured individually across three randomly selected fields. The proportion of damaged mitochondria (characterized by swelling, cristae disruption, or outer membrane rupture) was also calculated.

**TEM characterization of HNP1-NTP complexes.** For ultrastructural characterization of HNP1 in complex with the VDAC1 N-terminal peptide (NTP), 0.5 μM HNP1, 0.5 μM NTP, or an equimolar mixture of both were diluted in 20 mM Tris-HCl (pH 7.4), applied onto glow-discharged (15 s) 200-mesh formvar/carbon-coated grids (EMS) and incubated for 30 s, followed by negative staining with 2% uranyl acetate. Grids were examined using a Tecnai G2 Spirit transmission electron microscope (120 kV; Thermo Fisher Scientific, Hillsboro, OR, USA) at the Cryo-Electron Microscopy Center of Zhejiang University.

**Expression, purification and refolding of human VDAC1.** Expression, purification and refolding of recombinant human VDAC1 (hVDAC1) were performed as described previously with modifications<sup>5,6</sup>. The recombinant expression plasmid pET28(a)/hVDAC1, encoding hVDAC1 with C-terminal His tags, was transformed into *E. coli* BL21(DE3) competent cells. A single colony was inoculated into Luria–Bertani (LB) medium containing 50 μg/mL kanamycin and cultured at 37 °C until the optical density at 600 nm (OD<sub>600</sub>) reached 0.6. Protein expression was then induced by adding 1 mM isopropyl-β-D-thiogalactopyranoside (IPTG) and incubation was continued for approximately 4 h. Bacterial cells were harvested by centrifugation, resuspended in lysis buffer (50 mM Tris-HCl, pH 8.0, 2 mM EDTA, 584 mM

sucrose), and disrupted by ultrasonication. The resulting lysate was centrifuged, and the pellet was resuspended in wash buffer (20 mM Tris-HCl, pH 8.0, 2 mM CaCl<sub>2</sub>) and centrifuged again at 4 °C to collect inclusion bodies. The inclusion bodies were dissolved in denaturing buffer (20 mM Tris-HCl, pH 8.0, 100 mM NaCl, 6 M guanidine hydrochloride (GuHCl)) and then diluted with 20 mM Tris-HCl, pH 8.0, 100 mM NaCl to reduce the GuHCl concentration to 4.5 M. Imidazole was added to the resulting solution to a final concentration of 10 mM prior to purification. The solubilized inclusion bodies were subjected to purification using Ni-NTA affinity chromatography. The column was equilibrated with column buffer (20 mM Tris-HCl, pH 8.0, 100 mM NaCl, 4.5 M GdnHCl) containing 10 mM imidazole. After sample loading, the column was washed with column buffer containing 50 mM imidazole, and hVDAC1 was eluted with column buffer containing 150 mM imidazole. The eluate was supplemented with 2% lauryldimethylamine oxide (LDAO) to prepare for refolding. The eluate was transferred into a dialysis bag (molecular weight cut-off: 10 kDa) and dialyzed against refolding buffer (20 mM Tris-HCl, pH 8.0, 0.1% LDAO, 5 mM DTT) at 4 °C for 24 h with three changes of fresh buffer. The refolded hVDAC1 was characterized by liquid chromatography–mass spectrometry (LC–MS) and circular dichroism (CD) spectroscopy.

**EGS crosslinking assay.** To directly assess the effect of HNP1 on VDAC1 oligomerization, recombinant hVDAC1 (50 µg/mL) was incubated with or without HNP1 (50 nM) in 20 mM HEPES buffer (pH 8.3) at 25 °C for 15 min. Crosslinking was initiated by the addition of 100 μM ethylene glycol bis (succinimidyl succinate) (EGS; Sigma, E3257) and proceeded at 30 °C for 15 min, followed by quenching with 20 mM Tris-HCl (pH 8.3). Control groups included hVDAC1 alone and hVDAC1 with HNP1 but without EGS.

To investigate the role of HNP1 in VDAC1 oligomerization in hepatocytes, primary mouse hepatocytes were treated with 4 μM HNP1, with or without 20 μM VBIT-12, in the presence of 0.5 mM palmitic acid for 24 h. Cells were then harvested and resuspended in PBS containing 500 μM EGS, and incubated at 30 °C for 30 min. The crosslinking reaction was terminated by the addition of 20 mM Tris-HCl (pH 8.3). Control groups included untreated cells and cells treated with palmitic acid alone. The extent of VDAC1 multimerization was analyzed by immunoblotting using an anti-VDAC1 antibody.

**Immunoblot analysis.** Protein samples from the EGS crosslinking assay were separated by electrophoresis on 4–20% Bis-Tris PAGE gels (GenScript, M00928) using MOPS running buffer, and then transferred onto polyvinylidene difluoride (PVDF) membranes. Membranes were blocked with 5% non-fat milk in TBST for 1 h at room temperature and probed with an anti-human VDAC1/porin antibody (rabbit monoclonal, 1:1,000, Abcam, ab154856;) overnight at 4 °C. After three washes with PBST, membranes were incubated with the corresponding horseradish peroxidase (HRP)-conjugated secondary antibodies for 1 h at room temperature. Protein bands were visualized using Clarity Western ECL Substrate (Bio-Rad), and chemiluminescent signals were captured using a Tanon 4600 imaging system (Tanon Science & Technology, China).

**Enzyme-linked immunosorbent assay (ELISA).** Serum HNP1-3 levels were measured using a Human HNP1-3 ELISA Kit (Hycult Biotech, HK-317) according to the manufacturer's instructions, using the serum samples from 12-week HFD-fed WT and *HNP1<sup>TG</sup>* mice as described above.

**Fluorescence polarization (FP) assays.** FP assays were performed to analyze the binding affinity between HNP1 and its binding partners. FP values were measured using a Spark® multimode plate reader (Tecan) with excitation/emission wavelengths of 470/530 nm. For each assay, the analyte was serially diluted twofold in the respective assay buffer, mixed with an equal volume (100 μL) of final concentration of 10 nM FAM-labelled peptide in a 96-well plate, and incubated at room temperature for 1 h with gentle mixing. Negative and blank controls consisted of the FAM-labelled peptide alone and buffer alone, respectively.

The dissociation constant ($K_d$) was calculated by fitting the data to the following equation as previously described<sup>7,8</sup>:

$$F = F_0 + \left( \frac{F_C - F_0}{2[FLP]} \right) ([FLP] + [P]) + K_d - \sqrt{([FLP] + [P] + K_d)^2 - 4[FLP][P]}$$

where $F$, $F_0$ and $F_C$ represent the measured FP, the FP of free FAM-labelled peptide (FLP, negative control), and the FP of the FLP–protein complex, respectively; $[FLP]$ and $[P]$ denote the final concentrations of the corresponding FAM-labelled peptide and the titrated protein/peptide, respectively.

To analyze the binding affinity between HNP1 and VDAC1, FP assays were performed using N-terminally acetylated, FAM-labelled HNP1 peptide (FAM-A11K-HNP1: Ac-ACYCRIPACIKGERRYGTCIYQGRLWAFCC) as the FLP. Recombinant hVDAC1 was serially diluted twofold from 2 μM to 0.00390625 μM in assay buffer (20 mM Tris-HCl, pH 8.0, 0.1% LDAO). For determining the interaction between HNP1 and the NTP, FAM-labelled NTP (FAM-labelled NTP: MAVPPTYADLGKSARDVFTKGYGFGLK(FAM)-NH<sub>2</sub>) was used as the FLP. HNP1 was serially diluted twofold from 2 μM to 0.000244 μM in assay buffer (20 mM Tris-HCl, pH 8.0, 0.1% LDAO).

**Mitochondrial membrane potential assay.** Mitochondrial membrane potential was assessed using a tetramethylrhodamine ethyl ester (TMRE)-based Mitochondrial Membrane Potential Assay Kit (Beyotime,

C2001S) according to the manufacturer's instructions. Hepatocytes were treated with 4 μM HNP1 alone or in combination with 20 μM VBIT-12 under lipotoxic conditions for 24 h, followed by incubation with TMRE working solution (diluted from the supplied 1000× stock solution with assay buffer) and Hoechst 33342 (Beyotime, C1026) for 30 min at 37 $^{\circ}$C. Fluorescence images were acquired using a BZ-X810 fluorescence microscope (KEYENCE).

**Fluorescence colocalization of HNP1 and VDAC1 *in vitro*.** BNL CL.2 mouse hepatocytes were seeded on coverslips and cultured overnight. Cells were then incubated with 4 μM FAM-labelled HNP1 for 24 h, washed twice with PBS, and stained with Mito-Tracker Deep Red 633 (Beyotime, C1034) and Hoechst 33342 (Beyotime, C1026) for 30 min at 37 $^{\circ}$C according to the manufacturer's protocols. After washing with PBS, coverslips were mounted onto glass slides. Images were obtained by a BZ-X810 fluorescence microscope (KEYENCE).

**RNA extraction and real-time quantitative PCR.** Hepatocytes were harvested and homogenized in 500 μL of FreeZol reagent (Vazyme, R711-01) at 4 $^{\circ}$C. Total RNA was isolated according to the manufacturer's instructions and reverse-transcribed into cDNA using the HiScript III 1st Strand cDNA Synthesis Kit (+gDNA wiper) (Vazyme, R312-01). Real-time PCR was performed using 2× Phanta Ultra Master Mix (Vazyme, P518). The mRNA

expression levels of target genes were normalized to that of *β-actin* and calculated using the $2^{-\Delta\Delta C_t}$ method.

**2D NMR characterization.** For determination of interaction between HNP1 and NTP in different solvents, $^{15}\text{N}$-labelled NTP was diluted in an NMR buffer (10 mM PB [pH 6.8] and 10% $\text{D}_2\text{O}$) in the presence and absence of 40% TFE or 0.2% SDS to a working concentration of 400 $\mu\text{M}$, followed by titration of HNP1 from a stock solution to the NTP samples at a molar ratio of 0.2:1. For characterization of the dose-dependent influence of HNP1 on the 2D NMR spectra of NTP, $^{15}\text{N}$-labelled NTP was diluted in 10 mM PB (pH 6.8) supplemented with 10% $\text{D}_2\text{O}$ to a working concentration of 200 $\mu\text{M}$. HNP1 was then titrated from the stock solution to these NTP samples at molar ratios of 0:1, 0.5:1, 1:1, and 2:1, respectively. The NMR spectra were collected at 298.15 K on a Bruker Avance III HD 800 MHz spectrometer equipped with $^1\text{H}$-$^{13}\text{C}$-$^{15}\text{N}$ RT-probe. The operating frequencies were 800 MHz for $^1\text{H}$, 201 MHz for $^{13}\text{C}$, and 80 MHz for $^{15}\text{N}$. 2D [$^{15}\text{N}$, $^1\text{H}$] HSQC spectra were acquired by using the standard Bruker pulse sequence hsqcetfpf3gpsi2. The spectral widths were set to 13.9503 ppm ($\text{F}_2/{}^1\text{H}$) and 36.0000 ppm ($\text{F}_1/{}^{15}\text{N}$). A total of 1,024 complex data points was collected in the direct dimension ($\text{F}_2$), and 256 complex increments were collected in the indirect dimension ($\text{F}_1$). The spectrum was accumulated with 8 scans per increment. Data were collected and processed by Topspin 4.4.1, and analyzed by CCPNMR3.2.2. Assignment of residues was performed by using NMRtist, a cloud computing service for fully automated analysis of protein NMR spectra (https://nmrtist.org/)<sup>9</sup>.

**Molecular dynamics (MD) simulations.** The crystal structure of HNP1 dimer was retrieved from the Protein Data Bank under accession code of 3GNY<sup>10</sup>, and the monomeric chain A in this crystal dimer was employed in the MD simulations of the present study. The three-dimensional structural model of NTP was directly extracted from the crystal structure of hVDAC1 (PDB: 6G6U). Both a HNP1 molecule and a NTP molecule were randomly placed in a rectangular box to construct simulation system, which were assigned the force field parameters from the ff19SB using LEaP<sup>11</sup>. The simulation system was then solvated in a cuboid periodic boundary box of the transferable interatomic potential with three points model (TIP3P) water molecules, maintaining solvent layers of 12.0 Å between the solute surface and box edges, and were neutralized with sodium/chlorine counter ions.

MD simulations were conducted by using Amber20 with all covalent bonds involving hydrogen atoms constrained by SHAKE algorithm. Prior to production runs, minimization, heat, and equilibration were conducted by parallel computation with assignation of 64 AMD CPU cores. The entire system was minimized for 5000-step steepest descent algorithm and 5000-step conjugated gradient method respectively, with a harmonic force constant of 5 kcal mol<sup>-1</sup>Å<sup>-2</sup> applied on heavy atoms of the solutes. The system was further subjected to energy minimization by 5000-step steepest descent and 5000-step conjugated gradient minimizations, with harmonic restrain of Cα atoms by 5 kcal mol<sup>-1</sup>Å<sup>-2</sup>. Finally, the system was fully relaxed by 10000-step of steepest descent, followed by 10000-step conjugated gradient minimizations, without any restraints. After minimizations, the system was heated from 0 K to 300 K by using Langevin dynamics at a constant volume, with a time step of 2 fs and a harmonic force constant of 2 kcal mol<sup>-1</sup>Å<sup>-2</sup>. The heated system was optimized by a 200-ps density equilibration, and subsequently undergone two steps of NPT (T = 300 K, P = 1 atm) equilibration with or without backbone restrain for total 200 ps. Finally, these systems were respectively submitted to a 200-ns NPT (T = 300 K, P = 1 atm) production simulation with three replicates. All production simulations were conducted on a single Graphical Processing Unit (GPU) of NVIDIA GeForce RTX 5090 by using the Amber20 Compute Unified Device Architecture (CUDA) version of PMEMD (pmemd.cuda) with isotropic pressure scaling. Gibbs free energy change ($\Delta G$) between HNP1 and NTP was calculated by MMPBSA.py.MPI module using the MM/GBSA approach<sup>12</sup>, and further decomposed into the contribution of individual residue in both peptides.

**Cytotoxicity assay.** Hepatocyte cell lines (THLE-2 and BNL CL.2) were seeded in 96-well plates and treated with VBIT-12 or VBIT-12<sup>pro</sup> at concentrations ranging from 0.195 to 100 μM, prepared by twofold serial dilution, for 24 h. Subsequently, 10 μL of WST-1 reagent (Beyotime, C0035;) was added to each well and incubated for 1 h. Absorbance was measured at 450 nm ($OD_{450}$) using a microplate reader. Cell viability was calculated relative to untreated control wells.

**Dynamic light scattering (DLS) analysis.** Dynamic light scattering measurements were performed using a Malvern Zetasizer Nano (Malvern Panalytical). HNP1, NTP and their equimolar mixture were prepared in 10 mM phosphate buffer (pH 7.4) at concentrations of 0.1, 0.5, 2.5, or 12.5 μM HNP1, respectively, and filtered through a 0.22-μm PES membrane. After equilibrating for 120 seconds at room temperature, the relative intensity of scattered light was recorded. Because scattering intensity is highly sensitive to particle size, the raw intensity distribution is strongly biased toward larger particles. we therefore report number distributions, which were converted from intensity distributions through Mie theory with the manufacturer's software (Zetasizer Software 7.12).

**Circular dichroism (CD) spectroscopy.** CD spectrum of folded hVDAC1 was collected by a Jasco J-725 spectrophotometer (Jasco) using a 1 mm pathlength quartz tube. The spectrum was scanned under a nitrogen atmosphere, from 200 to 250 nm at a step of 0.5 nm, a scan speed of 100

nm/min, a response time of 1.0 s, and a scan bandwidth of 1.0 nm as well as a 2 mm path-length cuvette at 25 °C, with four scans averaged per time. The refolded hVDAC1 sample (20 µM) was measured in phosphate-buffered saline (PBS) containing 1% LDAO (pH 7.4). The secondary structure composition was estimated using the K2D3 web server (https://cbdm-01.zdv.uni-mainz.de/~andrade/k2d3/).

**Statistical Analysis.** Statistical significance was determined using two-tailed unpaired Student's *t* tests for two-group comparisons or one-way analysis of variance (ANOVA) for multi-group comparisons. *P* < 0.05 was considered statistically significant. Correlation was assessed by simple linear regression. All analyses were performed using Prism 10.0 software (GraphPad Software Inc., San Diego, CA, USA). Data are presented as mean ± SD or mean ± SEM as indicated in the figure legends.

**Figure S1. Amino acid sequences of the four human neutrophil α-defensins HNPs 1-4.** The conserved residues are colored, including six Cys residues (orange), one Arg (blue), one Glu (red), and one Gly (green).

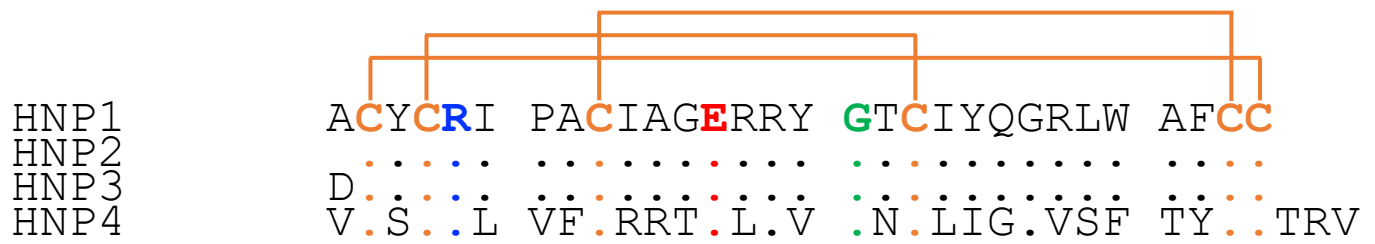

**Figure S2**

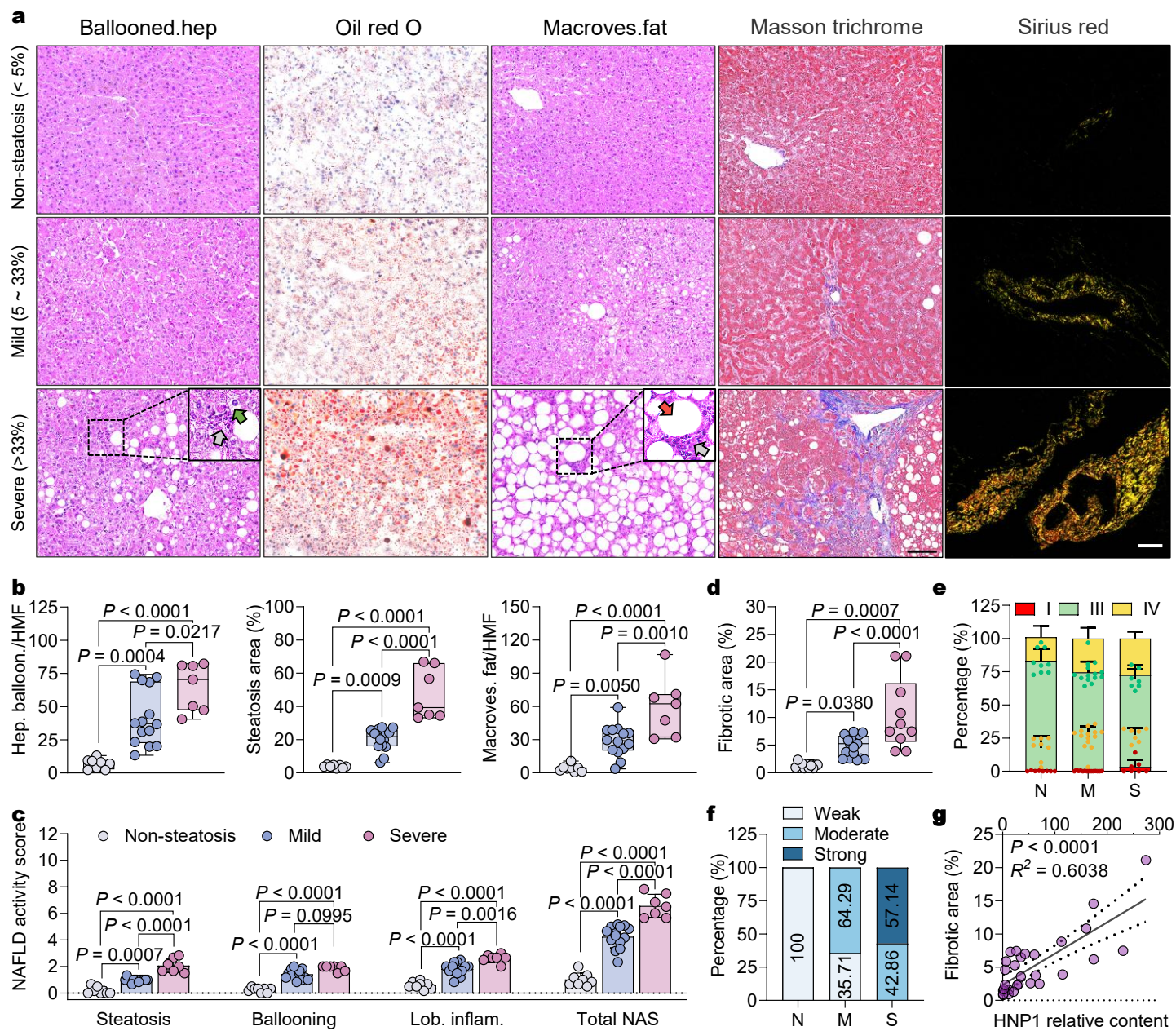

**Figure S2. Histopathological analysis of human liver samples.** (a) Representative images of human liver sections stained with H&E, ORO, Masson's trichrome, and Sirius red; gray, green, and red arrows indicate inflammatory infiltration, hepatocyte ballooning (Ballooned.hep), or macrovesicular fat (Macroves.fat), respectively. Scale bar: 200 μm. (b) Quantification of ballooned hepatocytes (left), ORO-positive area percentage (middle), and macrovesicular fat-laden hepatocytes (right) per HMF using ImageJ, based on three fields per section. (c) NAFLD activity score (NAS) of human livers without hepatic steatosis and with mild or severe steatosis. (d) Quantification of fibrotic area percentage in human livers based on Masson's trichrome-stained sections. (e) Percentages of different collagen types in Sirius red-stained human liver sections. (f) Percentages of human liver samples classified as weakly, moderately, or strongly positive based on IHC scoring. (g) Correlation analysis of hepatic HNP1 levels with fibrotic area. Coefficients of determination ($R^2$) and P values are shown. Statistical significance was calculated by one-way analysis of variance with the Bonferroni correction for multiple comparisons.

**Figure S3**

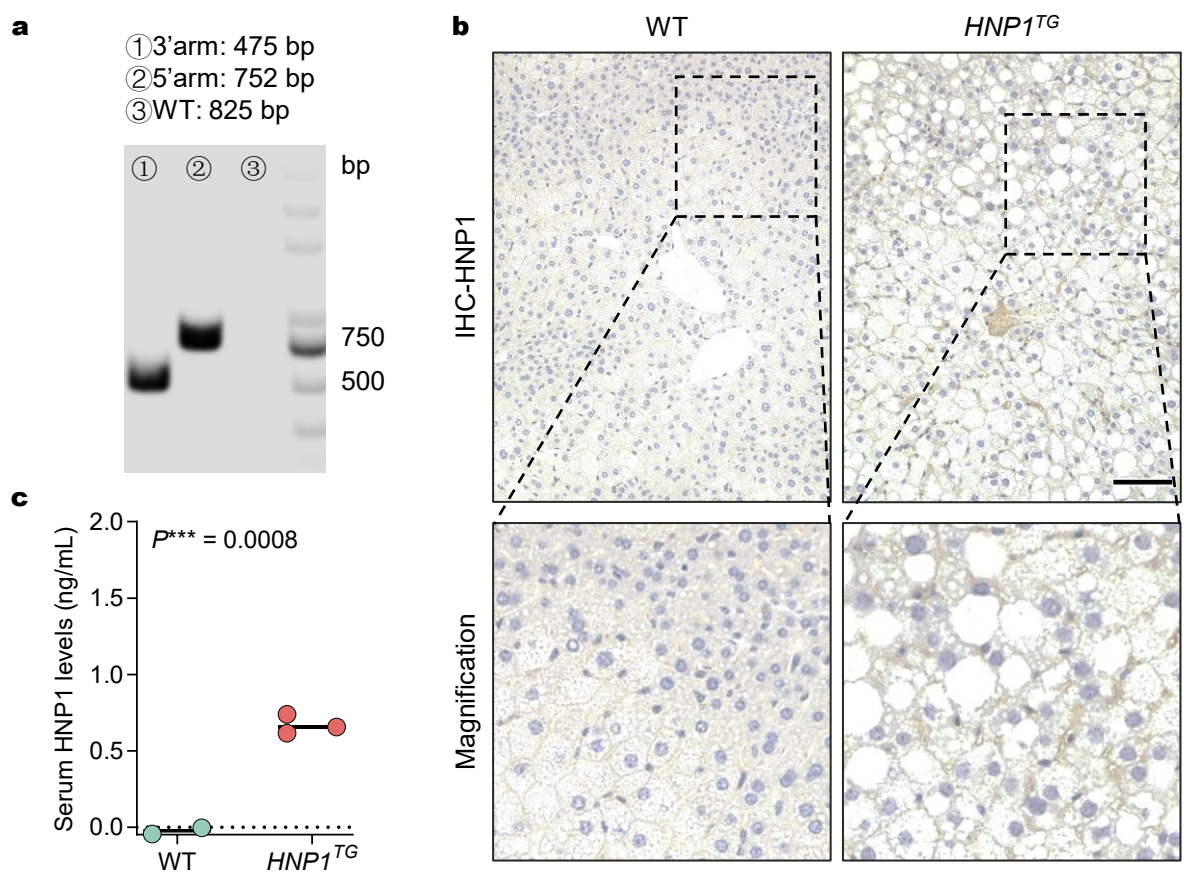

**Figure S3. Characterization of *HNP1<sup>TG</sup>* mice constructed by CRISPR-Cas9 using neutrophil elastase gene as a promoter.** (a) Successful knock in of DEFA1 in C57BL/6J mice characterized by semi-quantitative PCR. (b) Representative IHC images illustrating the levels and distribution of HNP1 in liver sections of WT or *HNP1<sup>TG</sup>* mice fed an HFD for 12 weeks. Scale bar, 200 μm. (c) HNP1 levels in serum from WT or *HNP1<sup>TG</sup>* mice fed an HFD for 12 weeks, determined by ELISA.

**Figure S4**

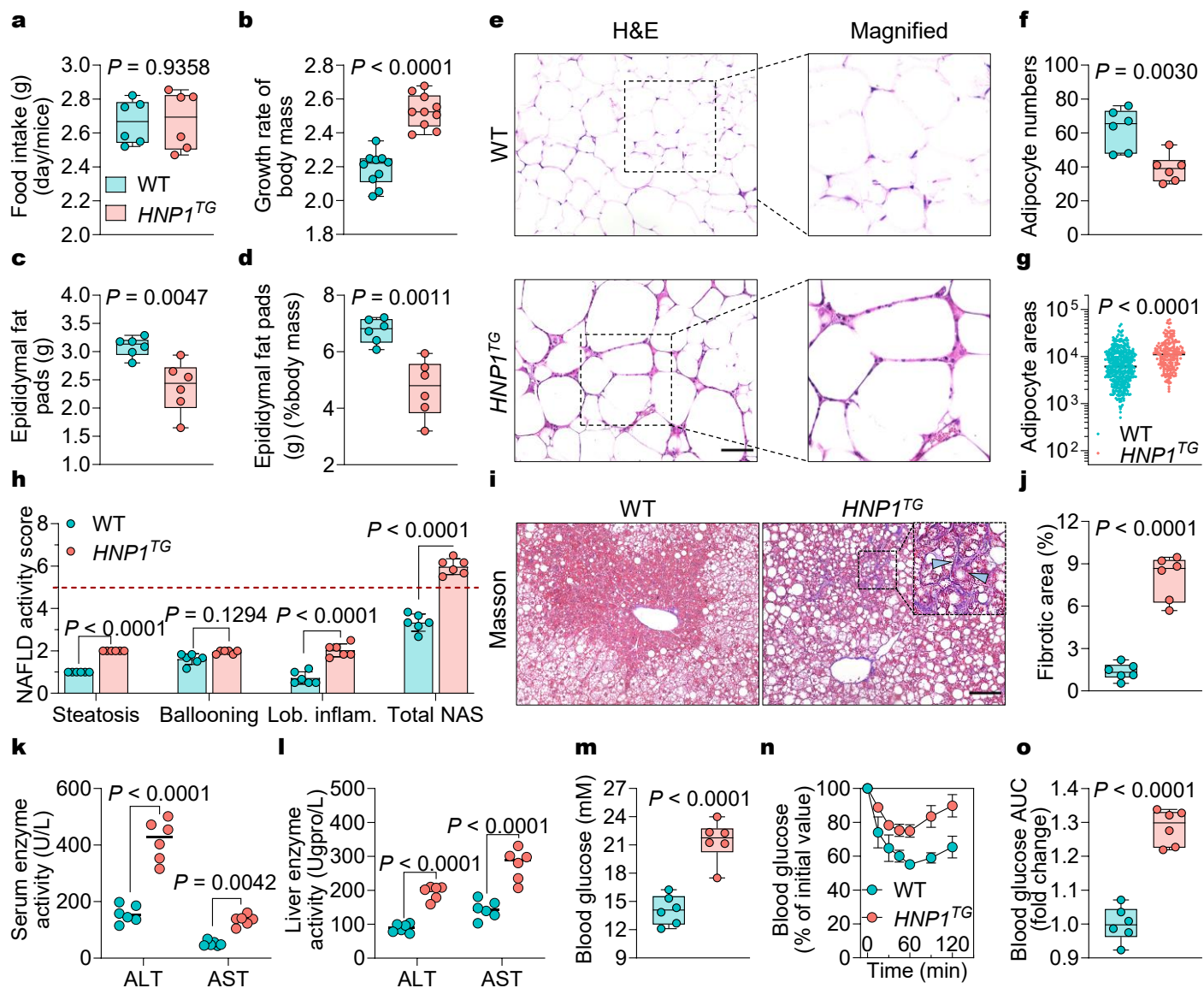

**Figure S4. Characterization of WT or *HNP1*<sup>TG</sup> mice fed an HFD for 12 weeks to induce MASLD.** (a) Determination of daily food intake per mouse ($n = 6$ mice per group). (b) Growth rate of body weights of WT and *HNP1*<sup>TG</sup> mice following HFD feeding for 12 weeks ($n = 10$ mice per group). (c-d) EAT weights (c) and EAT weight/body weight ratios (d) of WT and *HNP1*<sup>TG</sup> mice after 12 weeks of HFD feeding ($n = 6$ mice per group). (e) Representative images of H&E-stained mouse EAT sections. Scale bar: 200 μm. (f) Adipocyte number count in EAT sections using ImageJ, based on three fields per section ($n = 6$ mice per group). (g) Quantification of adipocyte area in H&E-stained EAT sections from WT or *HNP1*<sup>TG</sup> mice (405 and 203 adipocytes measured, respectively). (h) NAFLD activity score (NAS) of livers from WT or *HNP1*<sup>TG</sup> mice following 12 weeks of HFD feeding ($n = 6$ mice per group). (i) Representative images of Masson's trichrome-stained liver sections from WT and *HNP1*<sup>TG</sup> mice following 12 weeks of HFD feeding. Scale bar: 200 μm. (j) Quantification of fibrotic area percentage in mouse livers based on Masson's trichrome-stained sections ($n = 6$ mice per group). (k-l) Levels of ALT and AST in serum (k) and livers (l) from WT and *HNP1*<sup>TG</sup> mice fed an HFD for 12 weeks ($n = 6$ mice per group). (m) Levels of fasting blood glucose in serum of WT and *HNP1*<sup>TG</sup> mice after 12-week HFD feeding ($n = 6$ mice per group). (n-o) Insulin tolerance test of WT and *HNP1*<sup>TG</sup> mice following HFD feeding for 12 weeks (n), with fold changes in the area under curve (AUC) of blood glucose ($n = 6$ mice per group) (o). All data are shown as mean ± SD. Statistical significance was analyzed by an unpaired, 2-tailed Student's *t* test.

Figure S5

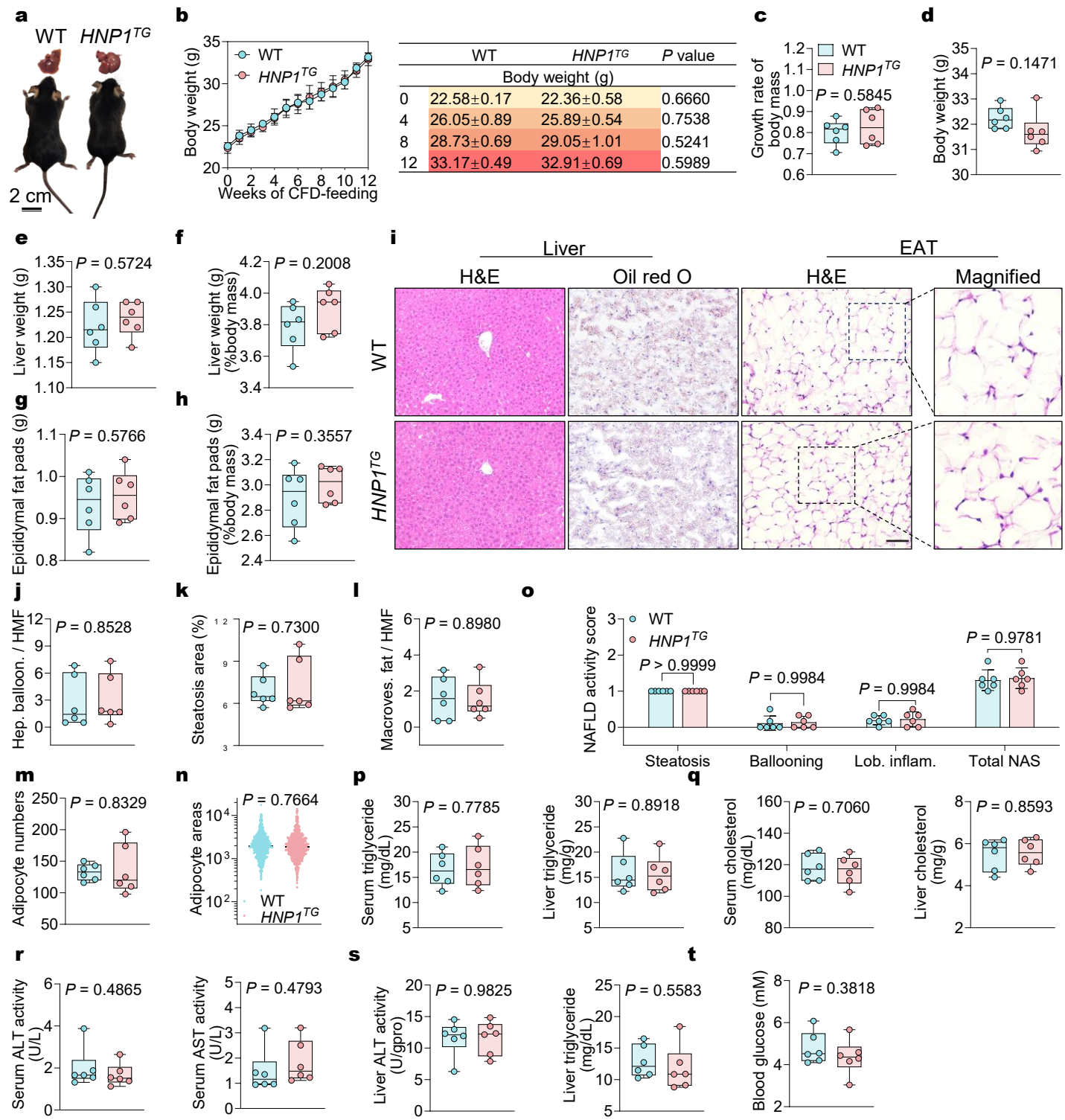

**Figure S5. Characterization of WT or *HNP1<sup>TG</sup>* mice fed a CFD for 12 weeks.** (a) Representative photographs of WT and *HNP1<sup>TG</sup>* mice (bottom) and their livers (top) following 12 weeks of CFD feeding. (b) Body weights of WT and *HNP1<sup>TG</sup>* mice measured weekly during CFD feeding ($n = 6$ mice per group) (left), and summary of body weights and statistical comparisons between WT and *HNP1<sup>TG</sup>* mice at 0, 4, 8, and 12 weeks (right). (c) Growth rate of body weights of WT and *HNP1<sup>TG</sup>* mice following CFD feeding for 12 weeks ($n = 6$ mice per group). (d-f) Body weights (d), liver weights (e), and liver weight/body weight ratios (f) of WT and *HNP1<sup>TG</sup>* mice after 12 weeks of CFD feeding ($n = 6$ mice per group). (g-h) EAT weights (g) and EAT weight/body weight ratios (h) of WT and *HNP1<sup>TG</sup>* mice after 12 weeks of CFD feeding ($n = 6$ mice per group). (i) Representative images of H&E- and ORO-stained mouse liver sections and H&E-stained mouse EAT sections. Scale bar: 200 $\mu\text{m}$. (j-l) Quantification of ballooned hepatocytes (j), percentage of ORO-positive area (k), and hepatocytes with macrovesicular fat (l) per HMF using ImageJ, based on three fields per section ($n = 6$ mice per group). (m) Adipocyte number count in EAT sections using ImageJ, based on three fields per section ($n = 6$ mice per group). (n) Quantification of adipocyte area in H&E-stained EAT sections from WT or *HNP1<sup>TG</sup>* mice (796 and 818 adipocytes measured, respectively). (o) NAFLD activity score (NAS) of livers from WT or *HNP1<sup>TG</sup>* mice following 12 weeks of CFD feeding ($n = 6$ mice per group). (p-q) Levels of triglyceride (p) and cholesterol (q) in serum and livers from mice after 12 weeks of CFD feeding ($n = 6$ mice per group). (r-s) Levels of ALT and AST in serum (r) and livers (s) from WT and *HNP1<sup>TG</sup>* mice fed a CFD for 12 weeks ($n = 6$ mice per group). (t) Levels of fasting blood glucose in serum of WT and *HNP1<sup>TG</sup>* mice after 12-week CFD feeding ($n = 6$ mice per group). All data are shown as mean ± SD. Statistical significance was analyzed by an unpaired, 2-tailed Student's $t$ test.

**Figure S6**

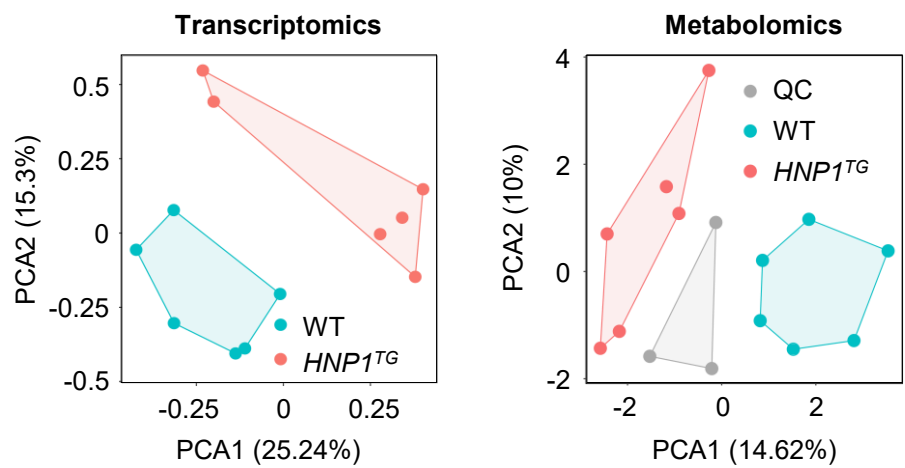

**Figure S6. PCA plots of transcriptomic and metabolomic data from WT and *HNP1<sup>TG</sup>* mice after 12 weeks of HFD feeding ($n = 6$ mice for each group).**

**Figure S7**

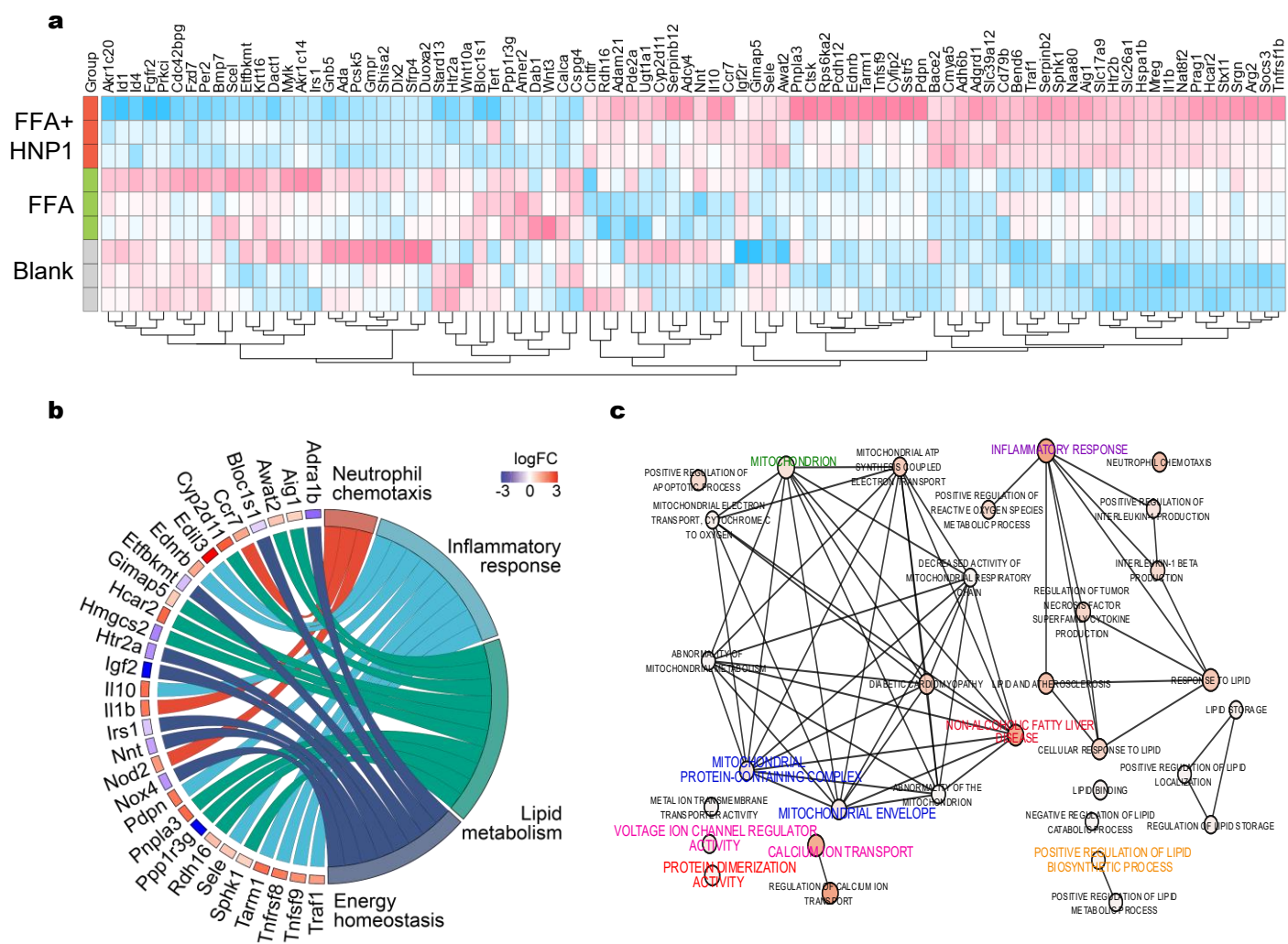

**Figure S7. Transcriptomic analysis of primary mouse hepatocytes treated with HNP1 in the presence of FFA.** (a) Heatmap showing expression patterns of immunometabolism-related DEGs in mouse hepatocytes under three conditions: blank (no FFA, no HNP1), FFA only, and FFA plus HNP1 ($n = 3$ for each group). The color key indicates the expression levels. (b) Circos plot showing several immunometabolism-related DEGs that are up- and down-regulated in hepatocytes treated with HNP1 in the presence of FFA, compared to those without HNP1. (c) Cytoscape network analysis of DEGs in hepatocytes treated with HNP1 in the presence of FFA, compared to those without HNP1.

**Figure S8**

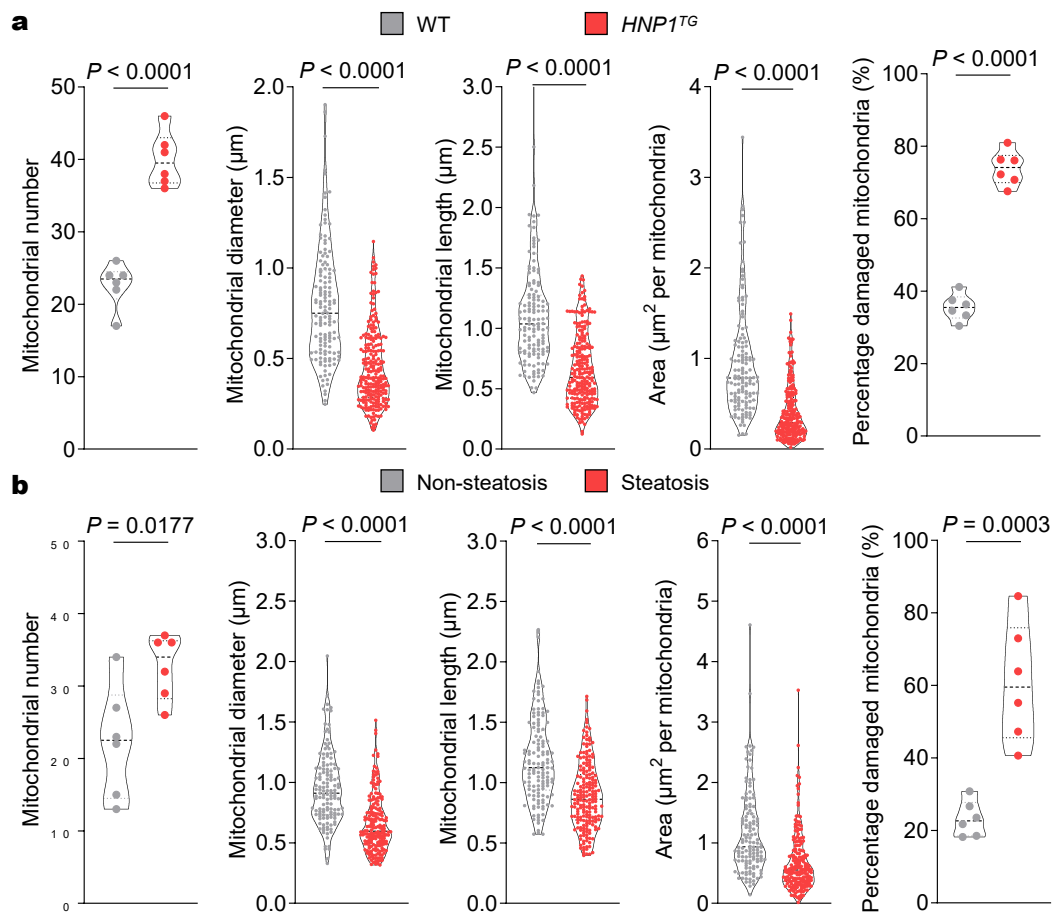

**Figure S8. Mitochondrial parameters of hepatocellular mitochondria in livers isolated from mouse models or clinical patients.** (a) Mitochondrial parameters of hepatocellular mitochondria in livers isolated from WT and *HNP1*<sup>TG</sup> mice post 12 weeks of HFD feeding. $n = 136$ mitochondria from 6 WT mice or 240 mitochondria from 6 *HNP1*<sup>TG</sup> mice. (b) Mitochondrial parameters of hepatocellular mitochondria in livers isolated from clinical patients with or without hepatic steatosis. $n = 196$ mitochondria from 6 patients with hepatic steatosis or 134 mitochondria from 6 controls. Data are shown as mean ± SD. Statistical significance was analyzed by an unpaired, 2-tailed Student's *t* test.

**Figure S9**

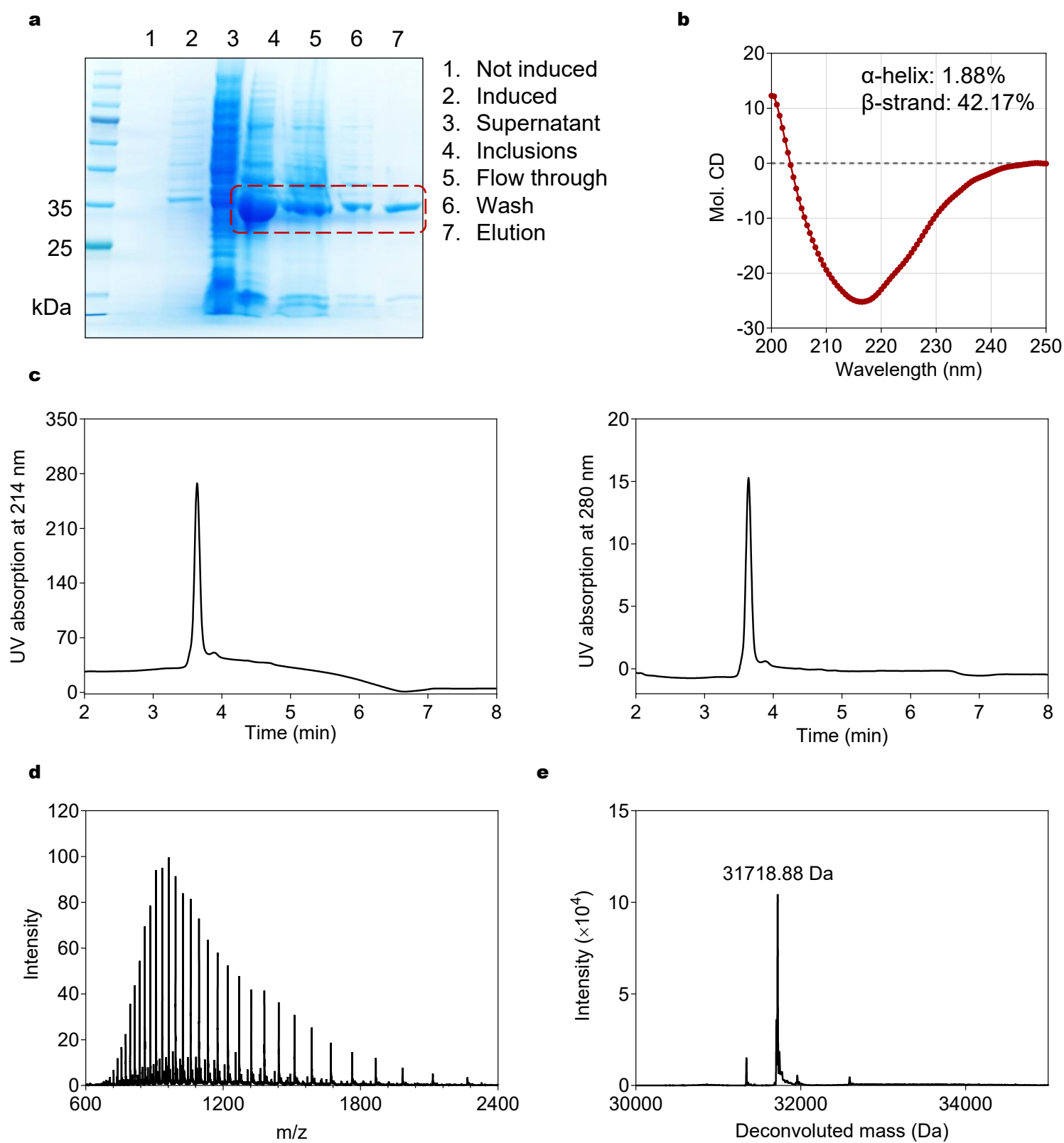

**Figure S9. Characterization of recombinant human VDAC1.** (a) SDS-PAGE gel analysis of the purified human VDAC1. (b) CD spectrum of the purified human VDAC1 refolded in 20 mM Tris-HCl (pH 7.4) containing 0.1% LDAO and 5 mM DTT. (c-d) The HPLC chromatograms (c) at 214 nm and 280 nm of the refolded human VDAC1 and its ESI-TOF mass spectrometric data (d). The chromatogram was obtained at 60 °C on an Agilent 300 SB-C18 Column (1.8 μm, 2.1 × 50 mm) running a solvent gradient of 5-65% B over 5 min at a flow rate of 0.2 mL/min (mobile phase A: 0.1% TFA in H<sub>2</sub>O, mobile phase B: 0.1% TFA in acetonitrile). (e) Deconvolution of the mass spectrum yields an observed molecular mass of 31718.88 Da, in agreement with the theoretical value of 31718.56 Da.

**Figure S10**

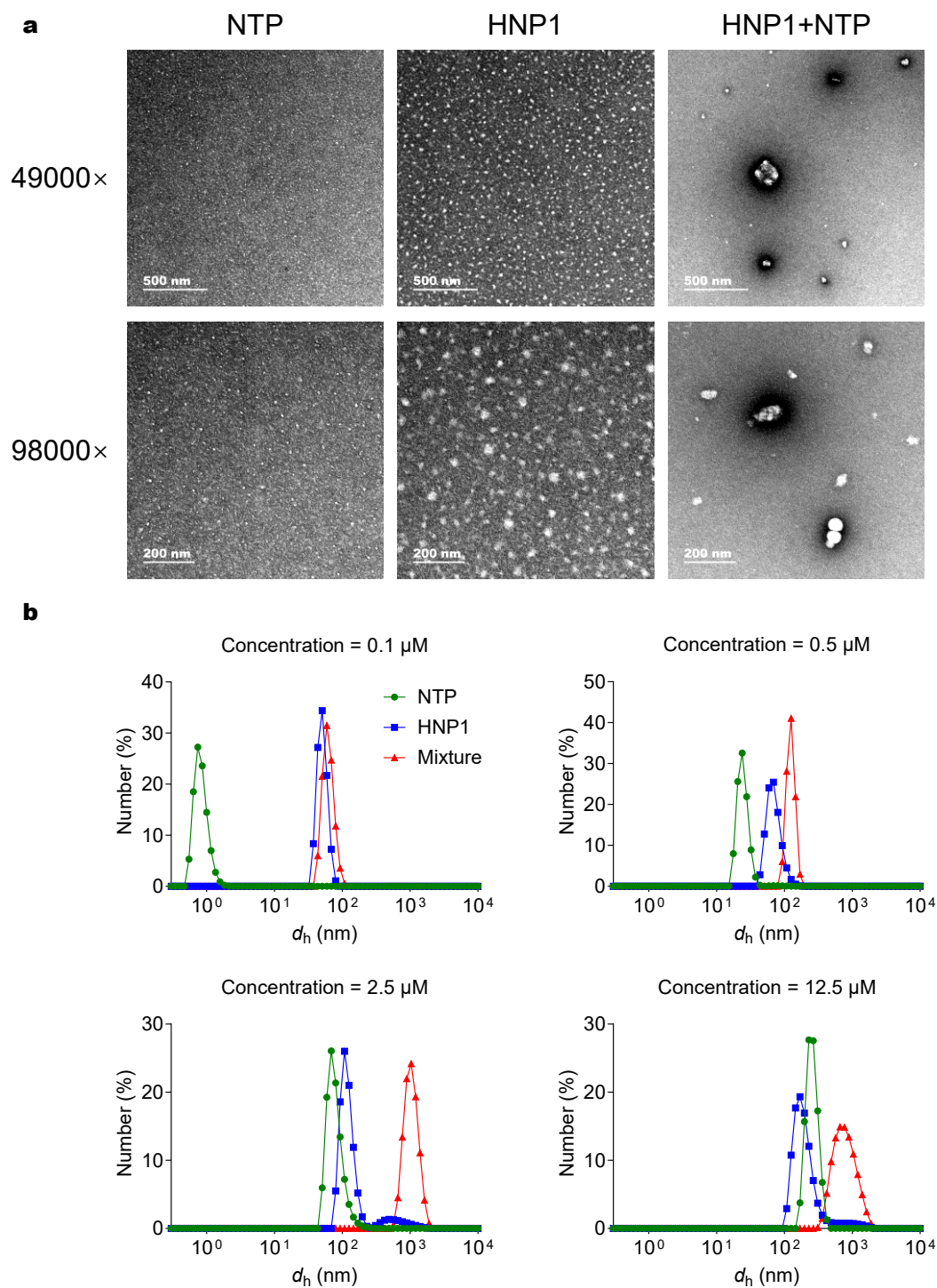

**Figure S10. Validation of HNP1 assembly with NTP in solution. (a)** TEM images of NTP (0.5 μM), HNP1 (0.5 μM), and their equimolar mixture in 20 mM Tris-HCl (pH 7.4). **(b)** Size ($d_h$) distributions of NTP, HNP1, and their equimolar mixture at peptide concentrations of 0.1 μM, 0.5 μM, 2.5 μM, or 12.5 μM, as determined by DLS.

**Figure S11**

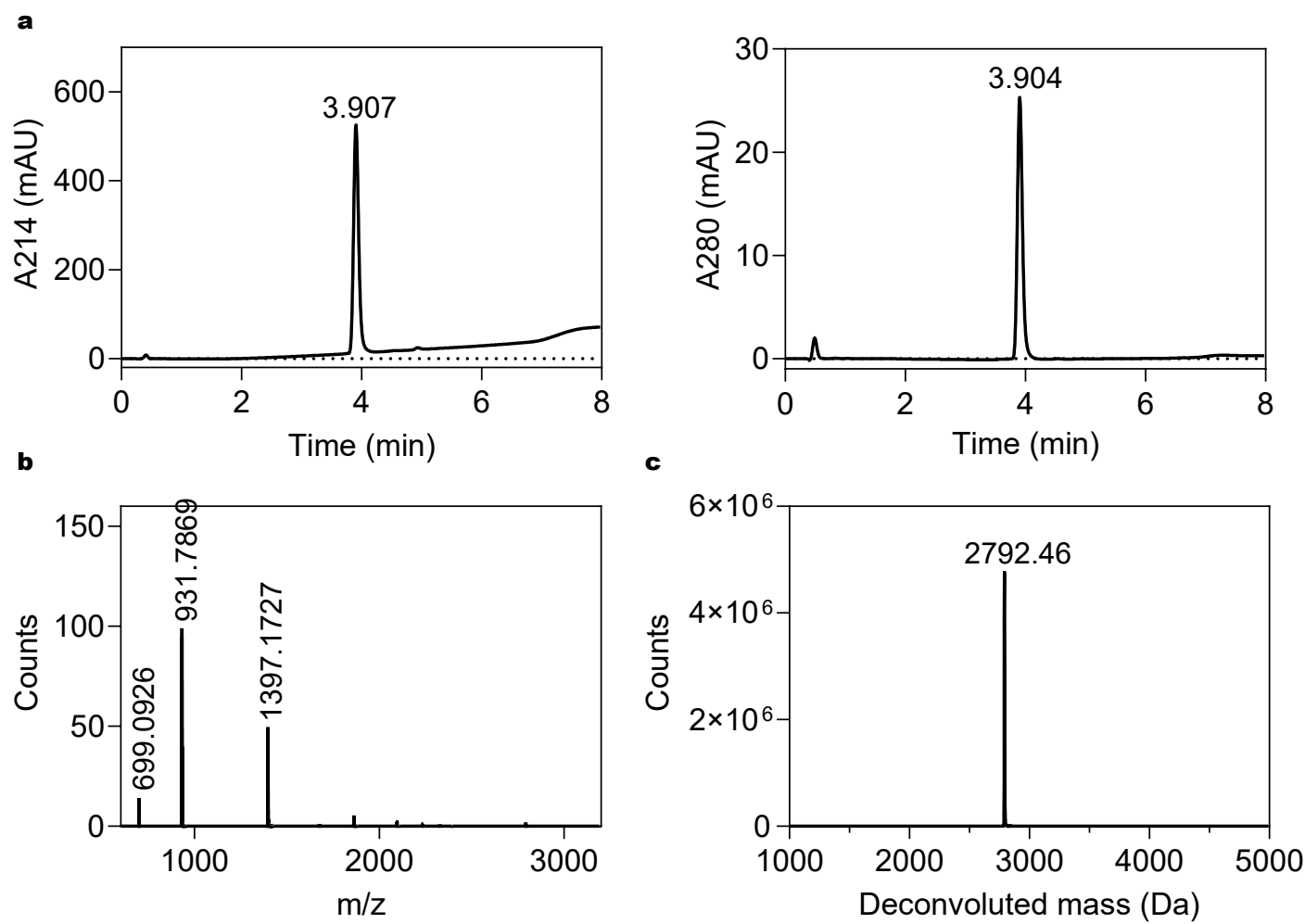

Calculated mass (wild-type NTP) = 2762.18 Da

Numbers of nitrogen atom = 31

Percentage of isotopic labeling = $(2792.46 - 2762.18) / 31 \times 100\% = 97.68\%$

**Figure S11. Characterization of the <sup>15</sup>N-labeled NTP peptide.** (a) The HPLC chromatograms of the <sup>15</sup>N-labeled NTP recorded at 214 nm and 280 nm. Chromatographic separation was performed on an Agilent 300 SB-C18 Column (1.8 μm, 2.1 × 50 mm) at 60 °C with a gradient of 5-65% solvent B over 5 min at a flow rate of 0.2 mL/min (mobile phase A: 0.1% TFA in H<sub>2</sub>O, mobile phase B: 0.1% TFA in acetonitrile). (b) ESI-TOF mass spectrum of the purified peptide. (c) Deconvolution of the mass spectrum revealed an observed molecular mass of 2792.46 Da, consistent with the theoretical value of 2793.18 Da, corresponding to an isotopic labeling efficiency of 97.68%.

**Figure S12**

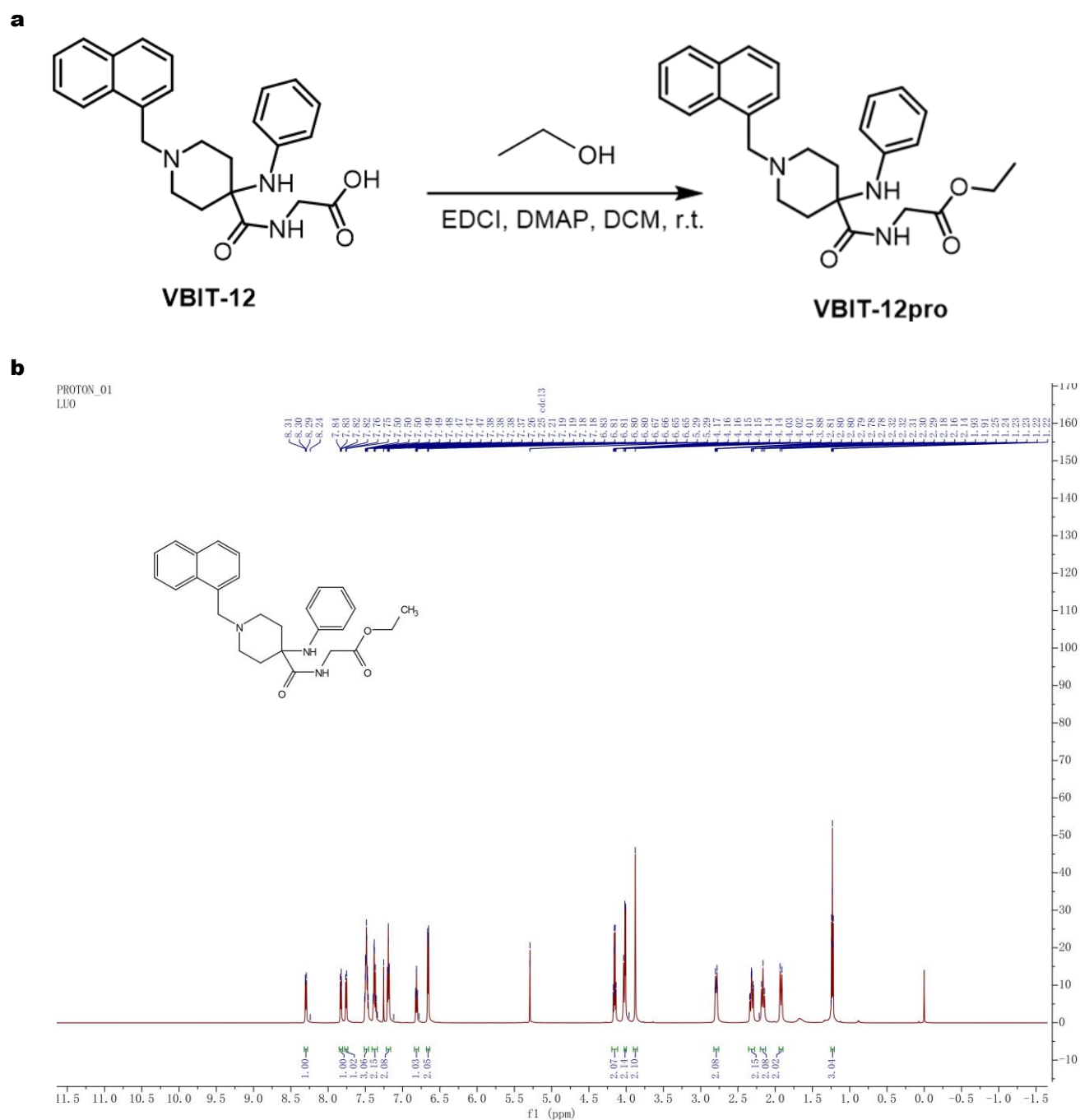

**Figure S12. Chemical synthesis of VBIT-12<sup>pro</sup>.** (a) One-step synthesis of VBIT-12<sup>pro</sup> from VBIT-12 via esterification. (b) <sup>1</sup>H NMR spectrum of VBIT-12<sup>pro</sup>.

Figure S13

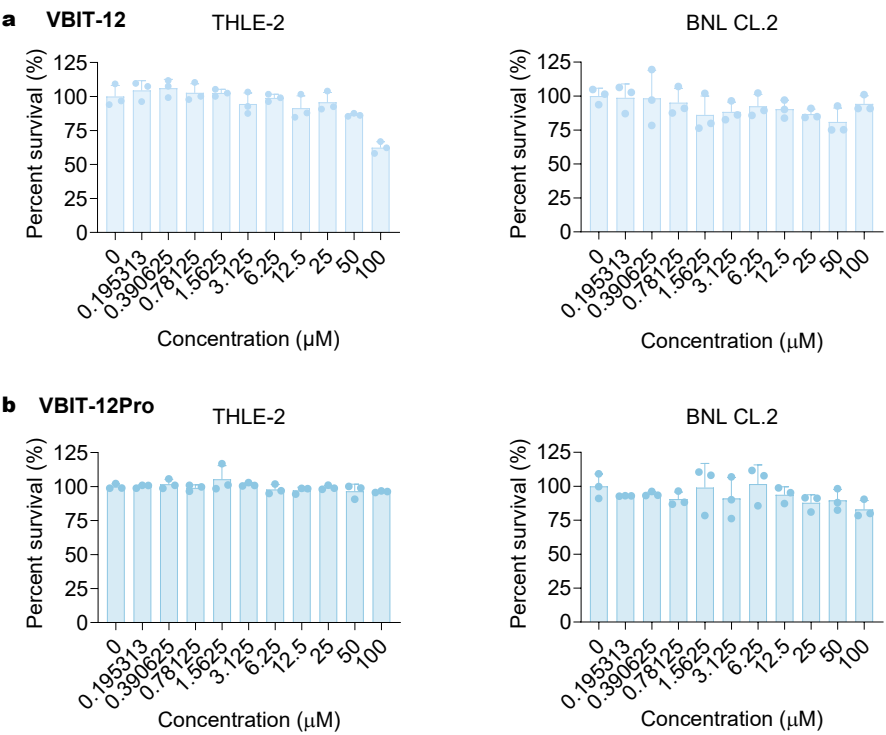

**Figure S13. *In vitro* cytotoxicity of VBIT-12 and VBIT-12<sup>pro</sup>. (a-b) *In vitro* cytotoxicity of VBIT-12 (a) and VBIT-12<sup>pro</sup> (b) to hepatocytic cell line THLE-2 and BNL CL.2. All data are shown as mean ± SD.**

Figure S14

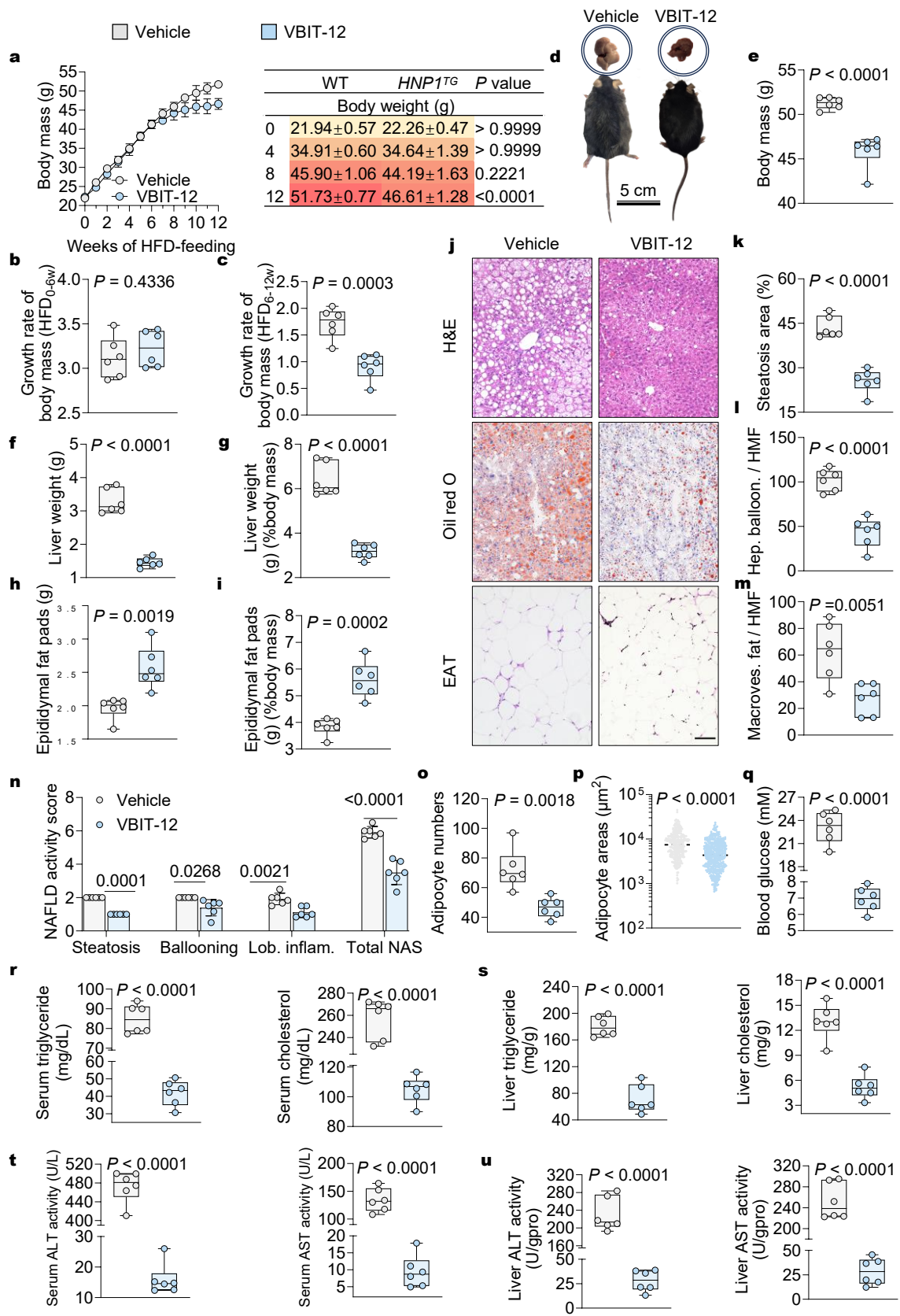

**Figure S14. Relieved MASLD in *HNP1<sup>TG</sup>* mice by intraperitoneal injection of VBIT-12.**

**(a)** Weekly body weights of HFD-fed *HNP1<sup>TG</sup>* mice treated with vehicle or VBIT-12 ($n = 6$ mice per group) (left), and summary of body weights and statistical comparisons between vehicle- and VBIT-12-treated *HNP1<sup>TG</sup>* mice at 0, 4, 8, and 12 weeks (right). **(b-c)** Body weight growth rates of vehicle- or VBIT-12-treated *HNP1<sup>TG</sup>* mice during the first 6 weeks **(b)** and the final 6 weeks **(c)** of a 12-week HFD feeding period ($n = 6$ mice per group). **(d)** Representative photographs of *HNP1<sup>TG</sup>* mice treated with vehicle or VBIT-12 (bottom) and their livers (top) following 12 weeks of HFD feeding. **(e-g)** Body weights **(e)**, liver weights **(f)**, and liver weight/body weight ratios **(g)** of vehicle- and VBIT-12-treated *HNP1<sup>TG</sup>* mice after 12 weeks of HFD feeding ($n = 6$ mice per group). **(h-i)** EAT weights **(h)** and EAT weight/body weight ratios **(i)** of vehicle- and VBIT-12-treated *HNP1<sup>TG</sup>* mice after 12 weeks of HFD feeding ($n = 6$ mice per group). **(j)** Representative images of H&E- and ORO-stained mouse liver sections and H&E-stained mouse EAT sections. Scale bar: 200 $\mu\text{m}$. **(k-m)** Quantification of percentage of ORO-positive area **(k)**, ballooned hepatocytes **(l)**, and hepatocytes with macrovesicular fat **(m)** per HMF using ImageJ, based on three fields per section ($n = 6$ mice per group). **(n)** NAFLD activity score (NAS) of livers from vehicle- or VBIT-12-treated *HNP1<sup>TG</sup>* mice after 12 weeks of HFD feeding ($n = 6$ mice per group). **(o)** Adipocyte number count in EAT sections using ImageJ, based on three fields per section ($n = 6$ mice per group). **(p)** Quantification of adipocyte area in H&E-stained EAT sections from vehicle- and VBIT-12-treated *HNP1<sup>TG</sup>* mice (279 and 435 adipocytes measured, respectively). **(q)** Levels of fasting blood glucose in serum from *HNP1<sup>TG</sup>* mice that were fed an HFD for 12 weeks and received daily treatment with vehicle or VBIT-12 for the last 6 weeks ($n = 6$ mice per group). **(r-s)** Levels of triglyceride and cholesterol in serum **(r)** and livers **(s)** from vehicle- and VBIT-12<sup>pro</sup>-treated *HNP1<sup>TG</sup>* mice after 12 weeks of HFD feeding ($n = 6$ mice per group). **(t-u)** Levels of ALT and AST in serum **(t)** and livers **(u)** from vehicle- or VBIT-12-treated *HNP1<sup>TG</sup>* mice fed an HFD for 12 weeks ($n = 6$ mice per group). All data are shown as mean ± SD. Statistical significance was analyzed by an unpaired, 2-tailed Student's $t$ test.

**Figure S15**

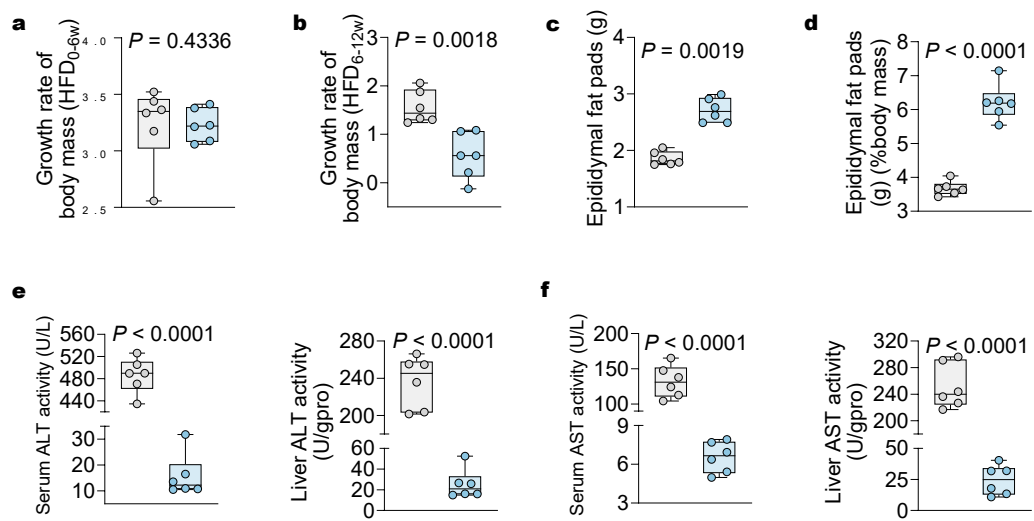

**Figure S15.** (a-b) Body weight growth rates of vehicle- or VBIT-12<sup>pro</sup>-treated *HNP1*<sup>TG</sup> mice during the first 6 weeks (a) and the final 6 weeks (b) of a 12-week HFD feeding period ($n = 6$ mice per group). (c-d) EAT weights (c) and EAT weight/body weight ratios (d) of vehicle- and VBIT-12<sup>pro</sup>-treated *HNP1*<sup>TG</sup> mice after 12 weeks of HFD feeding ($n = 6$ mice per group). (e-f) Levels of ALT and AST in serum (e) and livers (f) from vehicle- or VBIT-12<sup>pro</sup>-treated *HNP1*<sup>TG</sup> mice fed an HFD for 12 weeks ($n = 6$ mice per group). All data are shown as mean ± SD. Statistical significance was analyzed by an unpaired, 2-tailed Student's *t* test.
